# CyberSco.Py: open-source software for event-based, conditional microscopy

**DOI:** 10.1101/2022.03.16.484589

**Authors:** Lionel Chiron, Matthias LeBec, Céline Cordier, Sylvain Pouzet, Dimitrije Milunov, Alvaro Banderas, Jean-Marc Di Meglio, Benoit Sorre, Pascal Hersen

**Affiliations:** Institut Curie, Université PSL, Sorbonne Université, CNRS UMR168, Laboratoire Physico Chimie Curie, 75005 Paris, France; Laboratoire Matière et Systèmes Complexes, UMR 7057 CNRS & Université Paris Diderot, 10 rue Alice Domon et Léonie Duquet, 75013 Paris, France

**Keywords:** microscopy, automation, augmented microscopy, event-based microscopy, conditional microscopy, deep learning, image analysis, open-source software

## Abstract

Timelapse fluorescence microscopy imaging is routinely used in quantitative cell biology. However, microscopes could become much more powerful investigation systems if they were endowed with simple unsupervised decision-making algorithms to transform them into fully responsive and automated measurement devices. Here, we report CyberSco.Py, Python software for advanced automated timelapse experiments. We provide proof-of-principle of a user-friendly framework that increases the tunability and flexibility when setting up and running fluorescence timelapse microscopy experiments. Importantly, CyberSco.Py combines real-time image analysis with automation capability, which allows users to create conditional, event-based experiments in which the imaging acquisition parameters and the status of various devices can be changed automatically based on the image analysis. We exemplify the relevance of CyberSco.Py to cell biology using several use case experiments with budding yeast. We anticipate that CyberSco.Py could be used to address the growing need for smart microscopy systems to implement more informative quantitative cell biology experiments.

## Introduction

Microscopy imaging is an invaluable tool in quantitative cell biology. Recent years have seen the emergence of increasingly sophisticated techniques to probe the dynamics of living systems at high spatio-temporal resolutions. These technological developments have mostly been obtained using novel optical methods that structure the illumination of biological samples in space and time, the rise of optogenetics that facilitates real-time interactions with living samples, and the development of deep learning algorithms to analyze images and segment cells. Many microscopes are now powerful semi-automated systems that can acquire **pre-programmed timelapse sequences, usually via a process called <u>M</u>ulti-<u>D</u>imensional <u>A</u>cquisition (MDA), to observe and characterize the behaviors of single cells over extended periods of time**. To define a MDA protocol, the user typically has to select several locations within the biological sample (X and Y coordinates) and focal planes (Z positions), as well as the illumination settings that will be applied to every position (wavelengths, intensities, exposure time) and then, choose how often images should be captured by the camera. Automation microscopy software is used to ensure synchronization of the devices attached to the microscope, by periodically looping through these dimensions (space, time, imaging parameters). MDA has become very popular and is used routinely in cell biology laboratories. While the value of this approach has been well-demonstrated for the study of time-varying phenomena at play in biological systems, MDA drastically limits the capacity of fluorescence timelapse microscopy to monitor complex, multiscale biological processes. Indeed, for every experiment, a balance must be found between the number of positions imaged, the spatial resolution (magnification of a given objective), the time resolution, and additional effects such as phototoxicity, the duration of the experiment, cell density, etc. The tradeoffs between these factors are not trivial to setup and are usually not known at the beginning of experiments. Moreover, a simple MDA cannot typically deal with all of these factors; for example, studies of fast, intermittent processes (*e.g*., mitotic events) require imaging at both a high framerate and over a long period of observation, which lead to either phototoxicity or improper sampling.

Such basic workflows have become outdated at the time when smart systems and artificial intelligence are being used to improve the functioning of many (scientific) devices. A key practical limitation of MDAs is the fact that the images are only analyzed at the end of the experiment, which sequentially separates the workflows of image acquisition and image analysis. The ability to employ real-time image analysis to inform and optimize or adjust the settings of ongoing image acquisition would be a game changer for studying complex, dynamic cellular processes. Although this strategy requires a deep dive into the software programming and automation of microscopy devices, transformation of a conventional timelapse automated microscope into a powerful unsupervised automaton that is able to acquire data from a live biological sample at the right place and at the right timing could empower researchers in the biomedical sciences.

Building on existing automation microscopy software, several groups have started to explore how smart microscopy automate can benefit the life sciences^1–5^. In 2011, MicroPilot^5^ used LabVIEW© (a proprietary systems engineering automation software) to interface µManager (and other commercial vendor automation software) with a machine learning algorithm to identify and only focus on cells in a specific phase of mitosis. This strategy increased the throughput and decreased the time required to screen the desired cells. Since then, the rise of machine learning and the popularization of simple automation strategies—using Arduino, Raspberry and Python programming—have made it easier to build simple open-source solutions to achieve the same goals. For example, µMagellan^4^ and NanoJFluidics^1^ were built directly on µManager to achieve some level of feedback loop control and automation of image acquisition. µMagellan focuses on creating content-aware maps to adapt imaging modalities to a 3D biological sample. NanoJFluidics^1^ homemade array of Arduino-controlled syringe pumps combined with µManager can perform automated fixation, labeling and imaging of cells; notably, this system could be triggered by real-time detection of the rounding of mitotic cells through a basic image analysis algorithm. More recently, Pinkard et al.^3^ established a Python library that can interact with µManager to program a microscope in a very flexible way, though at the expense of a prerequisite for expert-level python coding skills. Overall, these examples harness the capability of µManager to pilot microscopy instruments and home-made image-analysis tool suites to trigger pre-programmed actions, and thus facilitate the development of complex or time-consuming microscopy experiments.

In addition, recent advances in the application of control theory to biology led to the development of external feedback loops, in which cells are analyzed and stimulated in real-time to force the cell state (*e.g*., expression of a gene^6–13^, activity of a signaling pathway^14^) to follow a user-defined (time varying) profile. Such feedback loops require the ability to perform real-time image analysis, in order to extract cellular features to feed an algorithm that decides how to stimulate the cells in live mode via microfluidics^10,11^ and/or optogenetics^6–9,15^. This novel and active field of research, called *cybergenetics*, harnesses the possibility of creating interactions between cells and a numerical model in real-time, and thus opens novel areas of both applied and fundamental research. To demonstrate the power of cybergenetics, we and others have developed various software to implement feedback loop-controlled microscopy systems. These solutions combine µManager and/or MATLAB with dedicated image analysis and control algorithms to close the feedback loop^10,11^. However, in practice, these solutions are difficult for non-experts to implement and cannot be easily transposed to a broad range of biological problems. More recently, several groups proposed Python and µManager-based software to develop cybergenetic experiments^16,17^, though these approaches remain specific to the control of gene expression in cells over time and do not meet all of the varied needs of cell biologists.

Here, we present CyberSco.Py software, which is a follow-up to our contributions to piloting gene expression in real-time in yeast and bacteria^10,11^. CyberSco.Py is written in Python, has been designed with automated, real-time feedback loops in mind, and includes deep learning image analysis methods as an integral part of image acquisition. We focused on achieving a proof-of-concept software with a simple, robust, user-friendly interface that can be deployed as a web application. Importantly, CyberSco.Py natively includes the ability to control basic microfluidic devices through an Arduino board that drives electro-fluidic valves (see Materials and Methods). CyberSco.Py is, by design, oriented towards advanced timelapse experiments that include triggered events and routed tree scenarios—rather than preprogrammed sequences of image acquisition. CyberSco.py is still a proof-of-concept and, here, our main goal is to demonstrate the potential of event-based, conditional microscopy to cell biologists. To this end, we first describe the principle of Cybersco.py, and then **focus on several use case scenarios to exemplify how event-based microscopy can be applied to perform more informative experiments relevant to cell biology**.

## Results

**CyberSco.py is an open-source web application for timelapse microscopy and microfluidics automation** written in Python (Figure 1). The software employs real-time image analysis and decision-making algorithms to trigger changes in the imaging parameters in real-time during the experiment. At present, this proof-of-concept is operational on a fully automated Olympus microscope (IX81) equipped with brightfield and epifluorescence illumination and linked to an Arduino-based homemade microfluidic control device (Supplementary Figure S1 and S2). A web interface allows the user to easily setup an experimental plan and/or to select pre-configured conditional experimental scenarios, together with the corresponding image analysis solutions. A local server receives these parameters and launches the experiment. During the experiment, the acquired images are constantly analyzed and used to trigger events according to the chosen scenario. In particular, the detected events can feedback on the microscopy settings and the microfluidic settings to adapt the experimental plan during the experiment (Figure 1). Thus, CyberSco.Py transforms microscopes and their related devices into an advanced imaging automaton capable of performing unsupervised time-dependent tasks with the capacity to handle various user-defined triggers. More details of the CyberSco.Py source code and its documentation are available on the GitHub page of the project (Supplementary Information). Although the software has been developed for a given microscopy setup, it can be extended to any equipment providing there is a driver and/or a documented communication protocol (see Supplementary Information and the GitHub page of the project on how to proceed).

**Figure 1.**
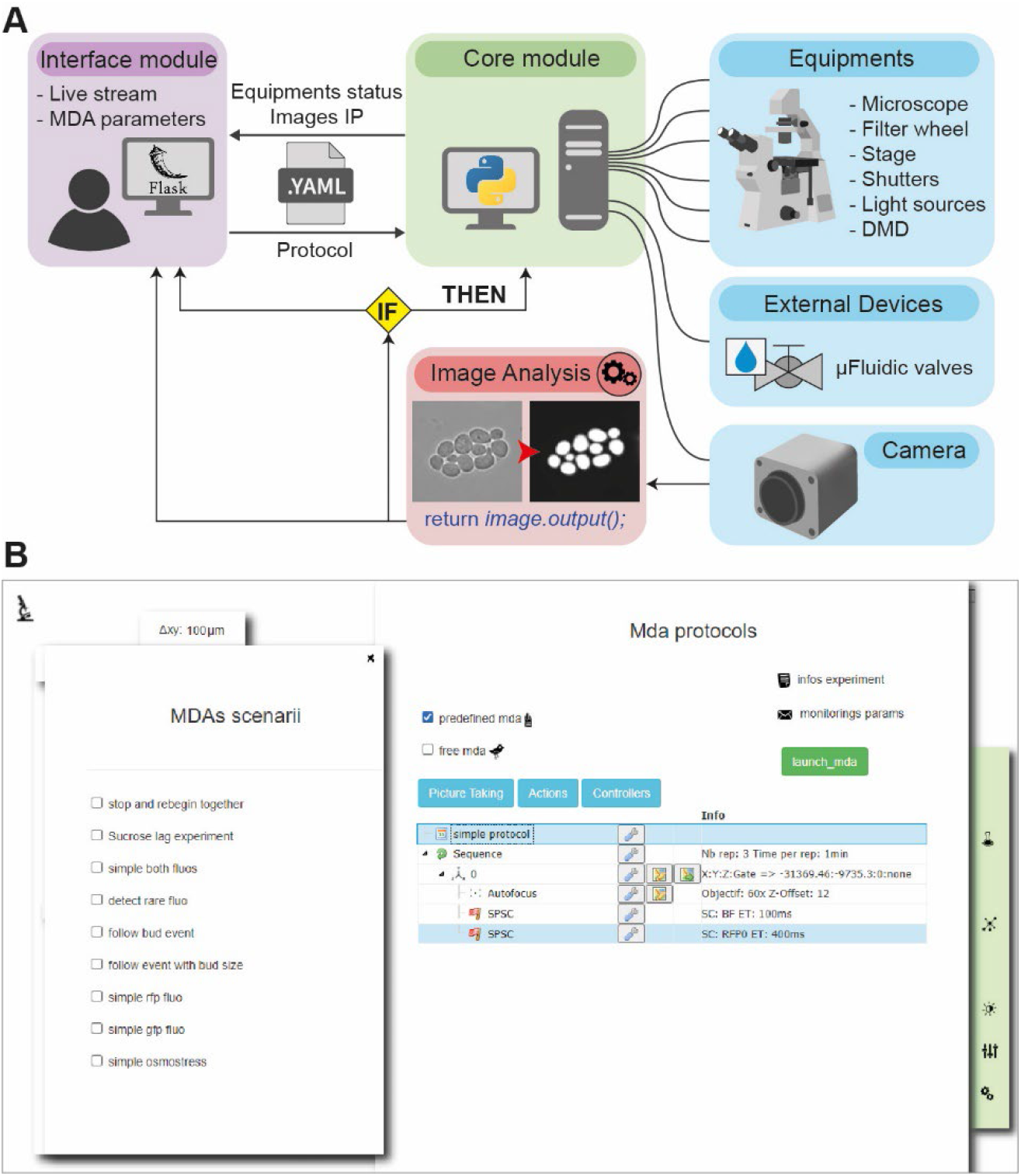
CyberSco.Py framework. **A**. Architecture. CyberSco.Py is built in Python and uses the web application library Flask to create a web user interface. Microscopy protocols are written into a YAML (human readable data serialization language) file, which can be interpreted by the Python core module of CyberSco.Py, which drives the various components of a IX81 fully automated microscope. The core module also drives a set of fluidic valves that can be used to switch the media flowing into a microfluidic device. A class in Python is associated to each device. Images obtained from the camera are analyzed in real-time by a U-NET deep learning model to segment yeast cells and/or detect specific events, depending on the pre-trained model selected by the user. The result of the analysis is used by the core module to update the current state of any devices under its control (See Materials and Methods for more information). **B**. Snapshot of the current user interface. The user interface is very simple by design and allows the user to choose between several pre-programmed event-based scenarios, for which the user must define various relevant parameters and condition switches. The simple drag and drop interface can be used to modify a given Multi-Dimensional Acquisition protocol to give more flexibility and to create more advanced protocols. The same interface can be used in “live mode” to view what is currently being imaged and check that the live image analysis is performing correctly. Once the program is launched, the computer takes control of the microscope and will adjust the image acquisition parameters based on the event-based scenario that has been selected. It is possible to code a novel scenario directly in Python and/or to manually adjust the thresholds and parameters used to detect events (*e.g*., number of cells, size of cells, etc.). The structure of a scenario consists of a list of instructions for the microscope (“make the autofocus”, “take a picture”, etc.) to be serially executed at each iteration, a conditional block, and an initialization block. Each scenario corresponds to a unique Python file with the same consistent structure. The user can also enter information about the projected experiment, as well as selecting modalities for monitoring the experiment remotely via email (selecting where to send the emails and at which frequency) and/or through a discussion channel (*e.g*., Microsoft Teams or Slack).

### From classic MDA to advanced MDA (aMDA)

CyberSco.Py can obviously be used to build classic MDA experiments. A web interface enables the imaging acquisition settings to be easily defined through a drag and drop interface (Figure 1 and Supplementary Information). The user can define X-Y positions and the corresponding focal planes and set the illumination parameters, as in conventional microscopy software. CyberSco.Py makes it easy to perform a classic MDA experiment that follows, for example, the proliferation of a population of yeast cells in a microfluidic device (Figure 2A). Crucially, the user can define specific imaging settings for each position (Figure 2B). This modification of how MDA is defined through the user interface is simple but powerful: by design, the user has full control over the acquisition settings without having to follow the classic MDA patterns of nested loops, which by default impose the same imaging acquisition parameters on all time points and positions. The ability to vary the imaging modalities per position imaged allows, for example, the user to conveniently and quickly optimize the imaging conditions by varying the exposure time for each position (to screen for phototoxicity or optimal illumination, for example). To demonstrate its usefulness, we used this feature to measure the light-dose response of a light-inducible promoter (Figure 2B and Supplementary Figure S3) with just one timelapse experiment. Setting up this experiment was quick and simple thanks to the minimal powerful user interface. More generally, any combination of imaging parameters can be assigned to a given position using the drag and drop tools within the user interface. For advanced users, the imaging parameters and positions can also be sent directly through a configuration file to create programmatically complex acquisition scenarios. More details of the user interface and scripting possibilities are available on the GitHub repository of the project (see also Supplementary Information). Notwithstanding such flexibility, CyberSco.Py has been programmed to include several types of protocols relevant to quantitative cell biology. Such built-in capabilities include: (1) synchronization of the image acquisition framerate with the microfluidic valve switches that apply the environmental changes; (2) detection and tracking of a cell of interest in a microfluidic device over an extended period of time; (3) cell counting, and triggering of environmental changes when the cell population reaches a certain size in the field of view; and (4) prediction of the future occurrence of a cellular event and the corresponding changes in the illumination settings and framerate required to image this event at an appropriate time interval.

**Figure 2.**
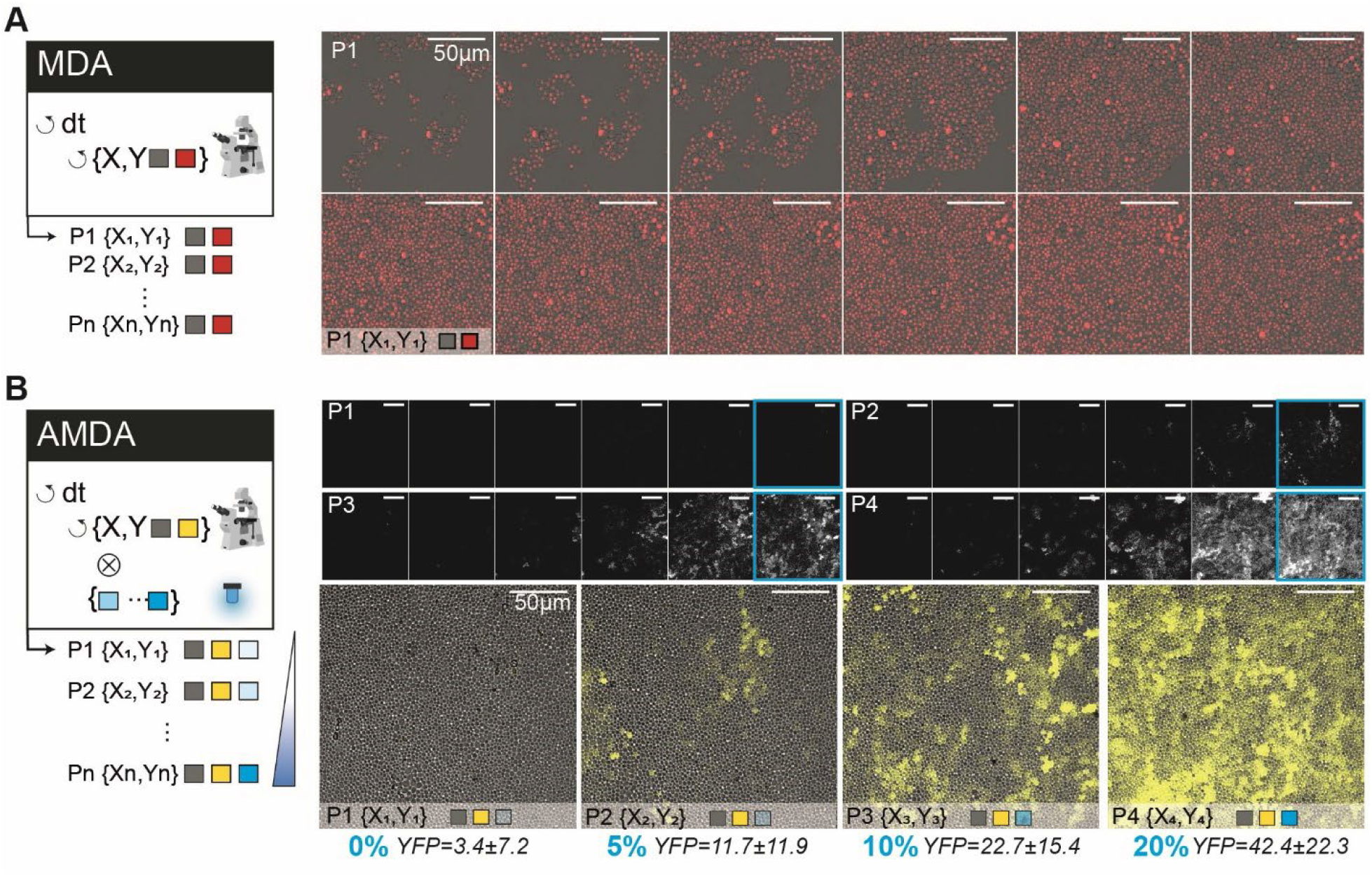
From simple to advanced MDA. **A**. Example of a classic Multi-Dimensional Acquisition (MDA) protocol to observe yeast proliferation in a microfluidic chamber, with two imaging channels (brightfield and RFP) imaged every 6 min for several hours. The HTB2 protein of the yeast cells is tagged with a mCherry fluorescent reporter. A sketch of the program (nested loops) is shown on the left side: the imaging parameters are identical for every position and timepoint. **B** An advanced MDA, in which the user has defined several positions, but set different illumination settings in the blue channel (LED intensity: 0%, 5%, 10% and 20%). This programming was done without scripts, by just using the drag and drop interface (see Supplementary Materials). Yeast cells bearing an optogenetic gene expression system (pC120-venus) were imaged for 15 hours. Each position is exposed to a different level of light stimulation, which alters the expression of a yellow fluorescent reporter both in terms of cell-cell variability, the maximum level of expression and dynamics. Thus, in one experiment, it was possible to quantitatively calibrate the pC120 optogenetic promoter using our settings without any requirement for coding (objective 20X). Fluorescence levels are averaged across the field of view and the error values are the standard deviation of pixel intensity.

### External triggers and adaptative acquisition framerates enable the observation of cell signaling at the right pace

Cells use a large set of signaling pathways and gene regulatory networks to process information from their surroundings. The signaling pathways in yeast are usually activated relatively quickly, within tens of seconds, while the transcriptional responses are slower (several minutes) and cell adaptation is even slower (tens of minutes). Therefore, it is difficult to image cell growth and signaling dynamics with fluorescence microscopy at the same time. Indeed, conventional MDA only allows image acquisition on one timescale. Fast periodic acquisition is possible, but leads to phototoxicity. Ideally, several acquisition frequencies need to be defined: a fast frequency to capture signaling events at the right pace, and slower frequencies, to image physiological adaptation and monitor cell growth. Moreover, the switch from a slow to fast acquisition framerate should be synchronized with the changes in the cellular environment through microfluidics. These technical requirements can be met by a simple scenario within Cybersco.py. As an example, we studied nuclear import of the MAPK Hog1p following hyperosmotic stress^18,19^. Yeast cells were grown inside a microfluidic device (See Materials and Methods) to facilitate imaging and facilitate dynamic environmental changes. We observed that an acquisition rate faster than one fluorescent image every 5-6 minutes led to phototoxicity and cellular arrest if performed over extended periods of time. This acquisition rate is too slow to capture nuclear localization of the Hog1p protein, which peaks one to two minutes after cells are subjected to hyperosmotic stress. We programmed Cybersco.Py to perform pulses of osmotic stress (by switching the state of an electrofluidic valve) every hour. Sending this command triggered modification of the acquisition framerate, which was increased from one frame every 6 minutes to one frame every 45 seconds (Figure 3). No coding/ scripting was required for this modification: the user just needed to select this predefined scenario and set the desired framerates and illumination parameters. As shown in Figure 3, we monitored several successive signaling events using this adaptative sampling rate without any user intervention. This example shows how the combination of external triggers and advanced MDA enables quantitative, time-resolved data on cellular responses and stress adaptation to be obtained without user supervision.

**Figure 3.**
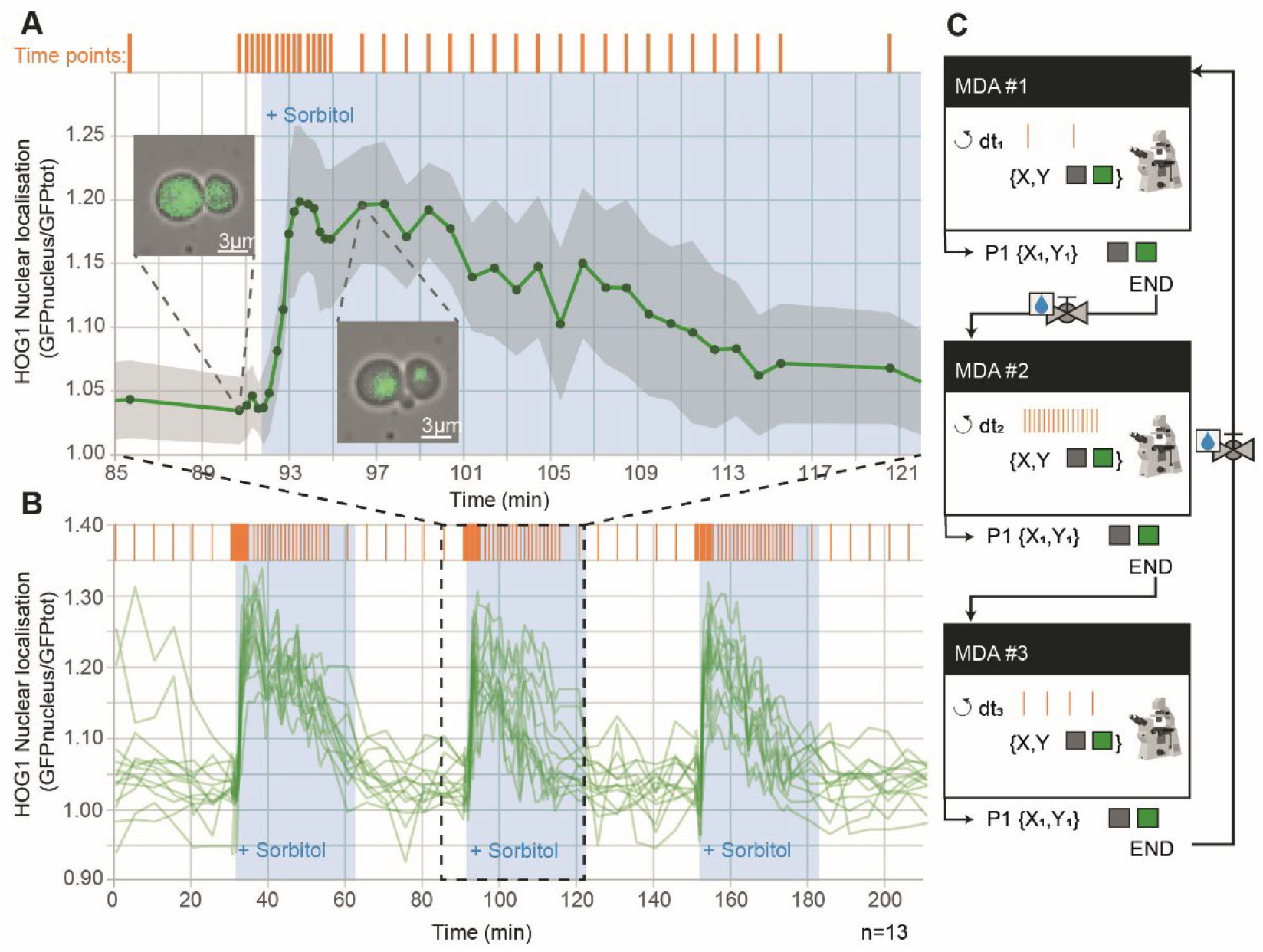
Synchronization of the acquisition framerate with dynamic perturbations to capture yeast cell signaling dynamics. **A**. Time course of nuclear accumulation of Hog1p in yeast cells growing as a monolayer in a microfluidic chamber subjected to an osmotic stress (1 M sorbitol). The insets show localization of Hog1-GFP before and after the osmotic stress. The acquisition framerate (orange bars) is automatically adjusted from one frame every 5 minutes to one frame every 15 seconds (20 times faster) just before the cells are stressed osmotically. The autofocus was turned off during the first 4 min of rapid Hog1 nuclear import. Recovery of the cells was then monitored at one frame every minute for 20 min, and finally the framerate was set back to its initial value (one frame every 5 minutes) until the next stress. The grey area represents the ± standard deviation of nuclear localization across 13 tracked cells from one microfluidic chamber. **B**. The adaptive sampling rate used in **A** was repeated three times to demonstrate that cells exhibit reproducible dynamics in response to every stress. This experiment allowed the timescales of activation (fast) and deactivation (slow) of the HOG cascade to be measured in an unsupervised manner. **C**. Sketch of the adaptive sampling MDA, which consists of three MDA experiments: one with a fast acquisition rate (nuclear import dynamics), one with a medium acquisition rate (nuclear export dynamics), and one with a slow acquisition rate (cell division after recovery). The switch from MDA#1 to MDA#2 is synchronized to activation of an electrofluidic valve that delivers an osmotic stress of 30 minutes duration (repeated every 60 minutes). Nuclear localization is computed as the mean of GFP fluorescence in the nucleus normalized to the mean of GFP fluorescence in the entire cell.

### Live cell segmentation enables the use of conditional events to dynamically change the modalities of image acquisition

CyberSco.py also offers the possibility of operating the microscope and the attached devices in real-time based on events detected during unsupervised analysis of the cell sample. The central idea is to let the microscope focus on “interesting” events through adjustment of the imaging acquisition parameters without supervision. This task requires efficient image analysis to segment cells, measure their properties and detect cellular events of interest. Image analysis is conveniently achieved in Python using the U-NET convolutional neural network^20^ (Materials and Methods and Supplementary Information), which is trained on a set of images. Once the training is complete, CyberSco.Py can use the resulting model to segment cells, display the segmentation in the user interface, compute cellular features and trigger user-defined events. At present, CyberSco.Py comes with two U-NET-trained models for yeast segmentation at different magnifications that give the following outputs: (1) the number of cells in the field of view; (2) a segmentation map of the cells in the field of view, as well as (3) their size and (4) their fluorescence levels. These cellular features can then be used to define conditional statements and adapt the imaging acquisition parameters in real-time. The segmentation results can be instantly visualized in “live” mode as a quality control step before launching timelapse experiments (see Supplementary Material). Below, we describe three different use case scenarios to exemplify the potential of conditional microscopy.

### Cells of interest can be detected and tracked in real-time

One interesting avenue of event-based microscopy is the ability to detect and focus on a particular cell of interest displaying a given phenotype at a given time. Instead of imaging many cells to find the cell of interest *a posteriori*, one can use real-time image analysis to identify cells with specific features and study their properties at an appropriate spatio-temporal resolution. There are two main challenges to overcome: defining the appropriate image analysis method to detect the cells of interest, and tracking those cells over time. Indeed, in an assembly of cells, growing cells push against their neighbors, often leading to large-scale displacement of the cells of interest, which may exit the field of view and be lost to subsequent imaging. Here, we demonstrate that it is possible to control the position of the X-Y stage to make sure that the cell of interest remains visible throughout the duration of the experiment. We studied a mixed population of yeast cells, in which a small fraction of the population (10%) express a fluorescent RFP histone tag. The program scans through the cells growing in a microfluidic device and once an RFP-expressing cell is detected, the scanning stops, the X-Y stage is moved to center the cell of interest and a timelapse is started to record RFP and brightfield images. The cell of interest is tracked throughout the timelapse, and the X-Y stage is moved so that this cell is always centered in the field of view. Figure 4 shows two such experiments, in different contexts, to demonstrate the efficiency of this detection and tracking scheme. Providing that the phenotype can be identified through image analysis (for example, a morphological feature or expression of a fluorescent reporter), this strategy could be employed to study rare phenotypes or, alternatively, to study long-term cellular behaviors (aging, cell-memory, habituation to repeated stress, etc.). within a large population of cells without user supervision.

**Figure 4.**
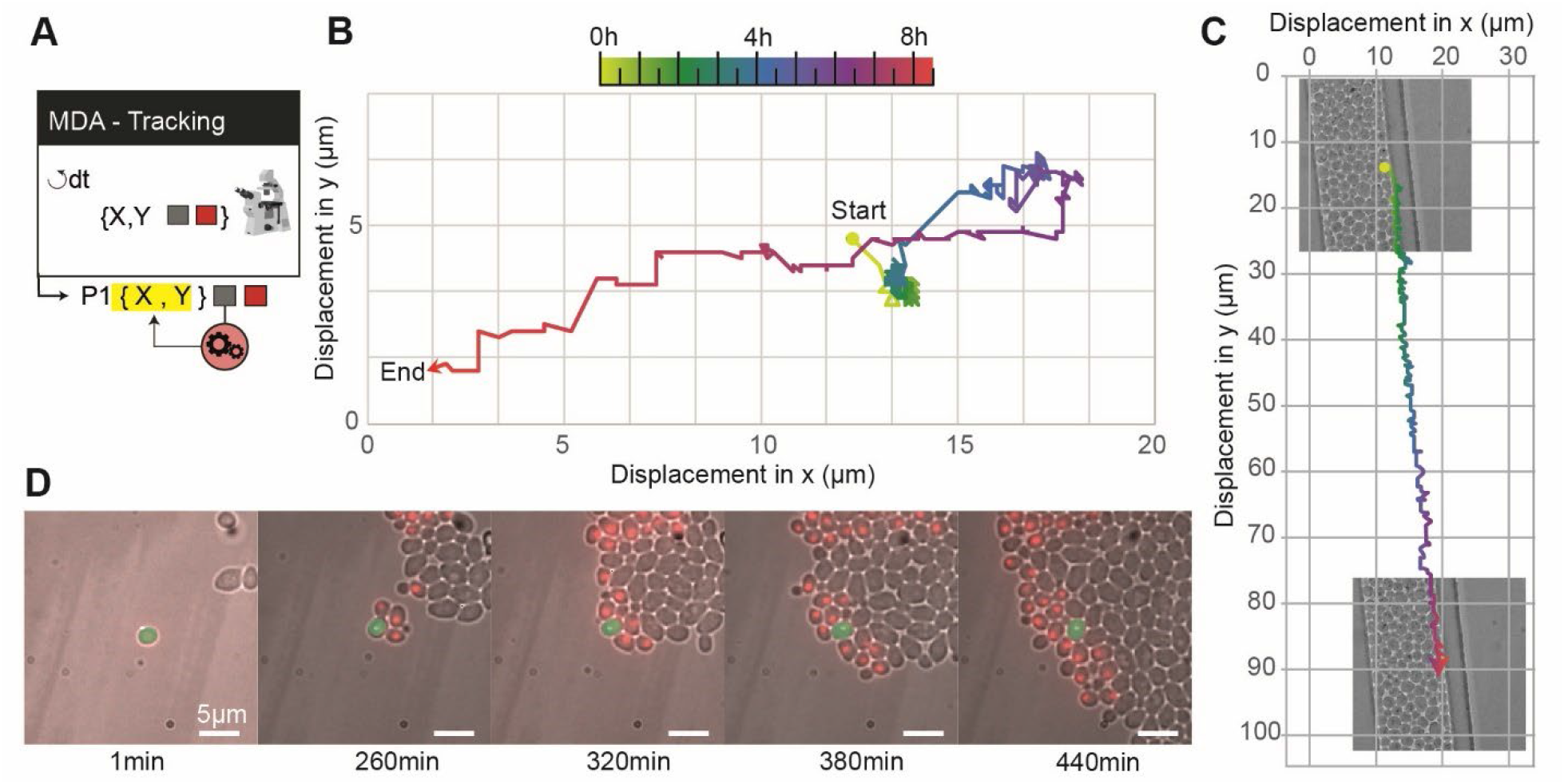
Detection and tracking of a cell of interest. **A**. Sketch of the “detect and track scenario”. Once a cell of interest is found in the field of view, the field of view is centered on that cell and the stage is periodically moved to maintain this cell in the center of the field of view. **B**. We mixed two populations of yeast cells in a microfluidic chamber, one of which express a HTB2-mCherry fluorescent reporter (1:10 cell ratio). The algorithm scans through several positions and when it detects cells with a signal in the RFP channel, picks one such cell randomly and centers it on the field of view. This cell is then tracked using brightfield segmentation, and the stage position is corrected through a feedback loop to compensate for cell displacement. **(C)** The cell of interest moves because it is pushed by the growth of neighboring cells, traveling approximately 20 µm during the course of the experiment. The real-time stage compensation keeps the cell in the center of the field of view. The duration of the experiment (around 9 h) is long enough to observe the appearance of the progeny of the cell of interest. **(D)** Tracking a non-fluorescent yeast cell growing in a dead-end narrow microfluidic chamber, leading to global directed motion of all cells. The tracked cell remains in the field of view, even though it travels approximately 80 µm; in contrast, the field of view is only ∼25 × 25 µm.

### Improving experimental reproducibility

Above, we showed how to control gene expression in cells based on real-time measurement of a fluorescent reporter. The same experimental strategy can be used to trigger a change in the cellular environment as a function of an observable feature in the field of view. This event-based strategy can be used to stimulate or perturb cells only when they have reached a given state. Alternative methods would require impractical, constant monitoring of the cells by the user. As a demonstration, we explored the impact of the number of yeast cells on the dynamics of recovery of cell division following a metabolic switch from glucose to sucrose. In response to glucose starvation, yeast cells produce and harbor the invertase Suc2p^21^ in their cell wall, which hydrolyzes sucrose into glucose and fructose in the extracellular environment. The yeast growth rate takes a certain amount of time to recover to normal after a metabolic shift from glucose to sucrose. Since the benefit of Suc2p production is shared among the yeast population, we hypothesized that the size of the population of cells may impact the response time after a metabolic shift to sucrose^21^. To test this assumption, cells growing in a microfluidic chamber were counted in real-time and, as soon as the number of cells in the chamber reached a given value (N=100, 500 or 2000; Figure 5), CyberSco.Py switched the perfusion from glucose to sucrose by triggering a microfluidic valve. Again, such experiments require an unsupervised, live method in order for the switch to be made efficiently. We observed that the duration of the lag phase decreased as the size of the population in the microfluidic chamber at the time of the metabolic shift increased, indicating faster production and accumulation of the enzymatic products within larger yeast populations, and hence a better adapatability of large yeast populations to sucrose metabolic shifts. Moreover, this experiment demonstrates the capacity of CyberSco.Py to precisely control the sample size at the start of the experiment, and suggests that cell density is a biologically relevant parameter that should be considered to improve experimental reproducibility.

**Figure 5.**
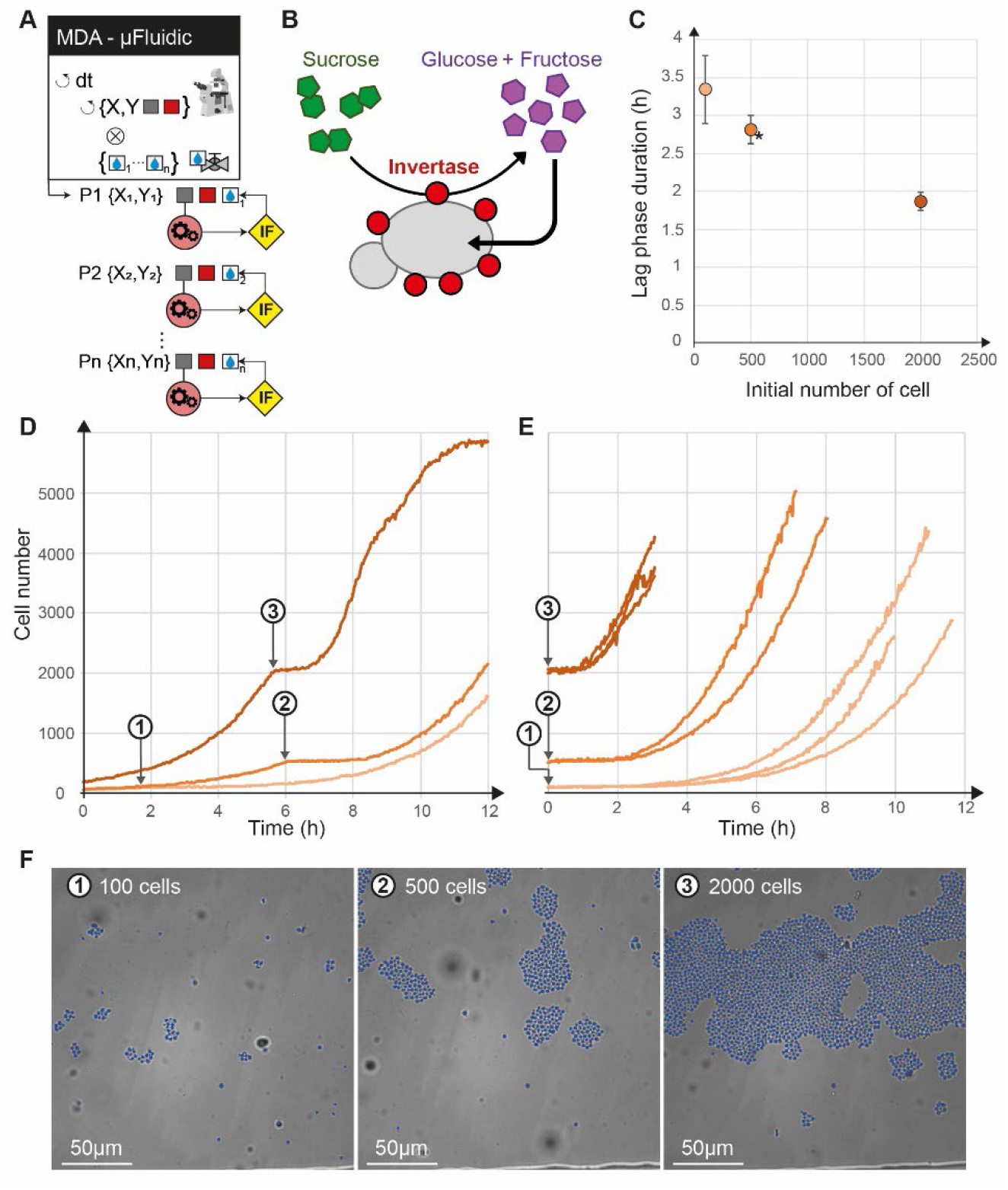
Conditional perturbation based on the number of cells. **A**. Sketch of the protocol, showing that different positions have a different conditional statement (<IF>) on the number of cells to trigger the switch from glucose to sucrose independently of each other. **B**. Sucrose conversion by yeast. The Suc2p invertase produced by cells is secreted extracellularly and can degrade extracellular sucrose into diffusible hexose. **C**. Following a shift from glucose to sucrose, cells need some time to convert sucrose to glucose and restart division. We show here that this time depends on the initial cell density (the higher the number of cells, the shorter the lag phase). The duration of the lag phase was estimated as the time it took the population to reach 130% of its initial size after the switch from glucose to sucrose. Error bars represent ± one standard deviation over three biological replicates (two replicates for the *). **D**. Temporal evolution of the number of cells for different initial densities: 100 (1), 500 (2) and 2000 (3) cells (grey arrows). **E**. Population growth shifted temporally to the switch time (i.e., switch = t_0_), demonstrating that the lag time increases as the initial cell density decreases. (F) Cell counting is achieved by real-time segmentation, shown here as an overlay of the brightfield image with single cell masks at the time of the valve switch.

### Switching between two MDAs by event triggering: imaging of mitosis in yeast at high temporal resolution

As a final example, we used CyberSco.Py to precisely image mitotic events in a population of growing yeast cells (Figure 6). We combined cell segmentation, cell tracking, and event-based modification of the imaging parameters to achieve imaging of mitotic events in yeast at a high temporal resolution in an unsupervised manner. Cells were observed and segmented at regular intervals (3 min). We assessed the increase in the size of buds over time to predict when mitosis will occur. We detected buds using U-NET segmentation and searched for yeast cells with buds that have grown to reach a threshold size and that have been increasing in size over the past three pictures (See Materials and Methods). Both criteria were sufficient to detect mitotic events under our conditions and to eliminate segmentation artefacts. When all conditions are fulfilled, the first MDA is stopped and a second MDA with a faster acquisition framerate, along with fluorescence imaging for the HTB2-mCherry reporter, is initiated. In this manner, mitotic events can be imaged at a much faster rate than in a classic MDA experiment (Figure 6) without the detrimental long term effect of phototoxicity. This kind of “search and zoom” scenario is presented for mitotic events as a proof-of-principle, but could be applied to any rare event that occurs within a population of cells.

**Figure 6.**
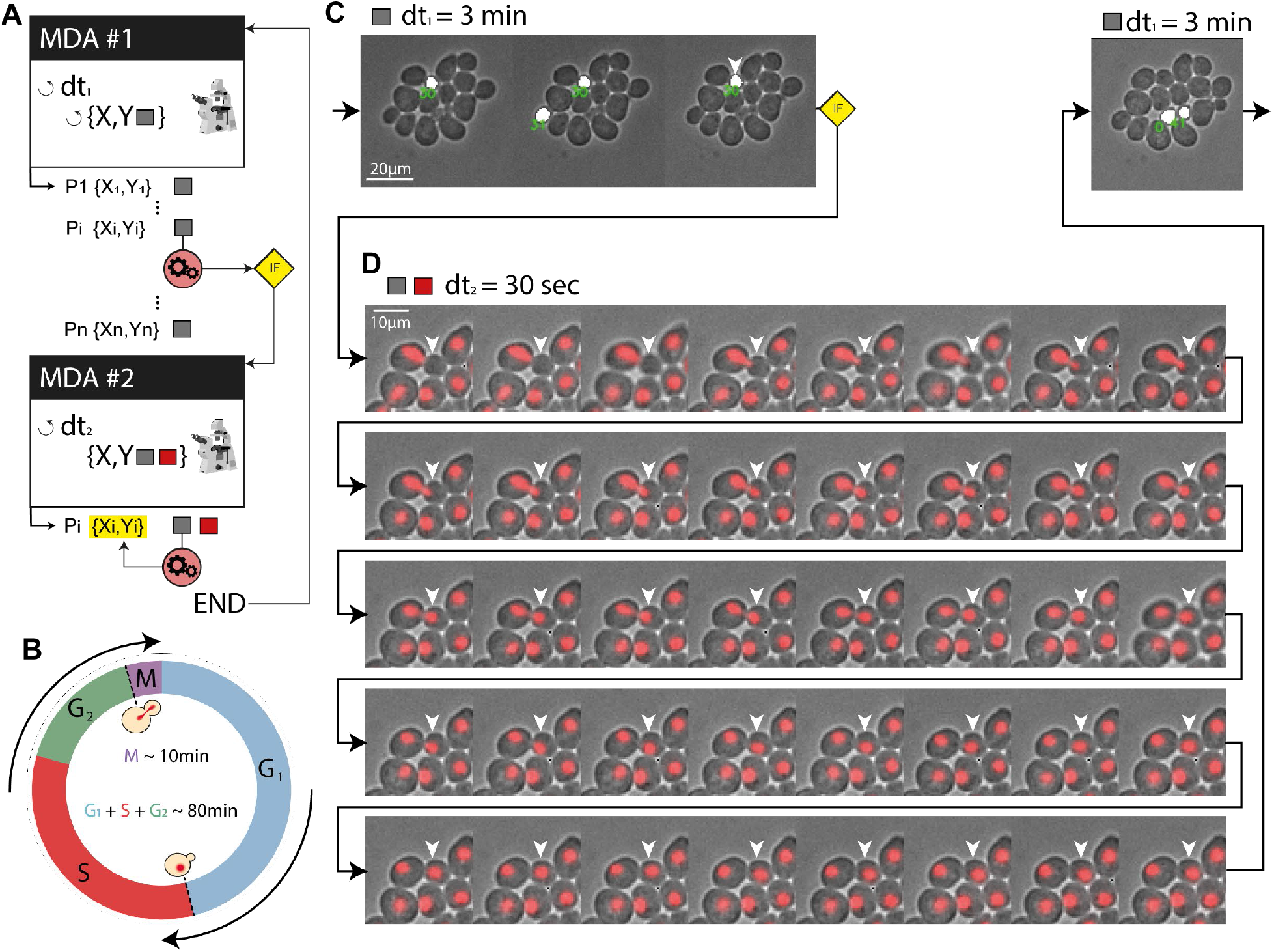
Bud detection and high-temporal-resolution imaging of mitosis. **A**. Scenario used to detect and “zoom in” on a particular event (in this case, mitosis). Several positions are monitored and when a condition is fulfilled, image acquisition is performed on only this position at an adapted sampling framerate. **B**. Cell cycle progression in yeast. The mitosis event to be captured represents a small-time fraction of the cell’s life cycle (∼10%). **C**. In practice, the acquisition of brightfield images of a population of budding yeast leads to a coarse timelapse with an acquisition framerate of 3 min to search for the next mitotic event. Cell segmentation is used to identify buds (size filtering), shown here as a white overlay. When a bud has reached a given size (and has been growing for at least three frames), we consider that a mitotic event is about to occur. **D**. Then, the acquisition software “zooms in” on that cell by increasing the framerate to one frame every 30 seconds for 20 minutes and RFP imaging is added to image the nucleus (HTB2-mCherry reporter). As shown in panel D, this scenario allows the complete mitotic event and nuclear separation between the mother and daughter cells (around 10 minutes, as expected) to be captured at an appropriate framerate. Once this image acquisition sequence is complete, the program resumes its search at the lower framerate for another mitotic event.

## Discussion

The main goal of CybserSco.Py is to enable the design of augmented MDA experiments in which the imaging settings can be changed in real-time as a function of unsupervised image analysis conducted during the time course of the experiment. CyberSco.Py has been built as a modular web application; in addition to the modularity of the device management, both the image analysis and decision-making algorithm are separate modules that can be adjusted individually and plugged into the communication modules that drive the microscope and its associated components. In that respect, CyberSco.Py aims to push forward “low code” or “no code” strategies, which are becoming increasingly popular, and enable biologists without coding expertise to create complex, event-based routines and workflows using cloud-based web applications. Creation of automation protocols that contain simple logical, conditional statements such as “IF this THEN that” or “WHILE this DO that” are within the reach of CyberSco.Py and may have an impact for researchers in biology who do not want, nor have the time or expertise, to dive into programming. This goal could be reached progressively by building on the examples proposed here and adding generic scenarios that are driven through the results of real-time image analysis. For example, the scenario developed in this work that acts on a microfluidic chip based on the number of cells could be directly reprogrammed by changing the trigger condition.

Image analysis remains the bottleneck of automation and will continue to require expertise in machine learning and coding. However, the pace at which deep learning is being adopted in laboratories and adapted into user-friendly software solutions and online tools^22,23^ suggests that an increasing number of easy-to-use methods to create UNET-trained networks will be developed in the near future and could subsequently be incorporated into CyberSco.Py. While we modified several classic experiments to demonstrate the potential of event-based microscopy, rethinking automation software for microscopy and including complete management of triggers and event detection may provide other important benefits in quantitative cell biology.

To start with, we can use our conditional microscopy framework to better prepare experiments and obtain a level of quality control before starting an experiment. Indeed, we showed that the number of cells may be an important factor when exploring the dynamics of population growth after a metabolic shift. Other processes related to cell-cell communication, metabolic gradients and cell-cell contact inhibition are also likely to be dependent on cell density. To increase experimental reproducibility, it seems reasonable to add a condition (or a set of conditions) on cell density to start a timelapse experiment. Similarly, only starting experiments, even simple timelapse studies, when a steady state is reached or when a gene has been expressed at a given level could improve experimental reproducibility. Only stimulating cells when the system is “ready” can also help avoiding phototoxicity and bleaching due to starting an experiment too early.

Timelapse experiments are usually long and prone to failures in both the microscopy system (*e.g*., loss of focus, improper control of temperature, drift in the stage) and the biological sample (*e.g*., contamination). However, in conventional timelapse microscopy, the user only realizes these issues at the end of the experiment, when it is too late. Our framework can be extended to regularly communicate the state of both the microscope and the experiment to the user. This could be in either a trigger mode, with the software sending a status report to the user through all sorts of classic communication channels (*e.g*. Slack, Teams, Email, SMS) when something goes wrong or the experiment reaches a given state, or—even better—the microscope could use the output of the image analysis to correct the problem automatically (*e.g*., by relaunching an autofocus step). Such simple automation workflows will certainly help to achieve high-quality data, reduce the time and cost of experiments, and improve experimental reproducibility.

Another important aspect is the huge amounts of data acquired in conventional, uninformed MDA. In many cases, this is due to the fact that image analysis is performed *a posteriori*, which requires as much data as possible to be acquired given the constraints of the imaging system and the biological sample. The ability to perform real-time analysis and conditional acquisition will make it possible to collect much sparser data, by focusing only on precisely what matters for a given study. This would speed up the analysis, facilitate data storage and sharing, and more generally improve the life cycle of imaging data.

We envision that advanced automation could be further used to perform online learning and automatically adjust the imaging parameters and stimulation of the biological system to obtain a model of the system under study. Such applications, which we are presently developing in the field of cancer research, may represent a game changer that increases the throughput of rare event detection and the quality of the resulting analysis by “zooming in” in time and space and/or sending drugs to perturb cells as soon as these rare events are detected among a large population of cells.

Our motivation to develop this proof-of-concept software with a simple user-friendly interface was to introduce conditional microscopy to a large audience. While this first step is relatively limited, the initial framework we propose here can be improved and developed further. Ideally, a global effort to develop application programming interfaces (APIs) for lab automation would facilitate the development of no-code workflows that are accessible to all researchers and integrate with commonly used collaborative online tools such as Slack and Teams. We believe that researchers could benefit from such advanced ways of conducting experiments, especially the ability to perform event-based automated imaging. CyberSco.Py is a first step to bring such concepts to the attention of biologists.

## Materials and Methods

### Yeast strains and growth conditions

All yeast strains used in this study are derived from the BY4741 background (EUROSCARF Y00000). The list of strains can be found in Supplementary Table 1. Yeast cells were picked from a colony in an agar plate, grown overnight in 2 mL YPD media, then the culture was diluted 1/100 in 5 mL of filtered synthetic complete media (SC; 6.7 g Yeast Nitrogen Base w/o amino acid [Difco 291940] and 0.8 g complete supplement mixture drop-out [Formedium DCS0019) to 1 L] supplemented with 2% glucose and cultured for 4-6 h at 30 °C with orbital shaking at 250 RPM (Innova 4230 incubator). The media used to perfuse the microfluidic chips during the experiments was SC supplemented with either 2% glucose or 1% sucrose. The microfluidic chips were made following a previously published protocol^11^ (see Figure S8). Liquid perfusion was performed using an Ismatec IPC (ISM932D) peristaltic pump at 50 µL/min (or 120 µL/min for the osmotic shock experiment). A homemade Arduino-based system was used to switch the state of an electrofluidic valve to change the media that perfuses the microfluidic chip. The microfluidic chip (Supplementary Figure S2) allows yeast cells growing in a monolayer to be imaged and has been described in previous works^24^. Another microfluidic design^25,26^ was used (in Figure 4E) to constrain cell displacement in one direction.

### Microscopy imaging

We used a fully automated Olympus IX81 inverted epifluorescence microscope equipped with a motorized stage (Prior Pro Scan III), Photometrix Evolve512 camera, and a pE-4000 CoolLed as a fluorescent light source. The objectives used in this study were either a 20X UPlanSApo or 60X PlanApo N. For the RFP channel, we used the 550 nm LED through a filter cube (EX) 545 nm/30; (EM) 620 nm/60 (U-N49005) with a 150 ms exposure time. For the GFP channel, we used the 460 nm LED through a filter cube (EX) 545 nm/30; (EM) 620 nm/60 (U-N49005) with a 150 ms exposure time. For the YFP channel, we used the 525 nm LED through a filter cube (EX) 514 nm/10; (EM) 545 nm/40 (49905 – ET) with a 500 ms exposure time. Microscopy experiments were carried out in a thermostat chamber set to 30 °C.

### CyberSco.Py software

CyberSco.Py is written in Python for the backend and HTML/CSS/ JS for the frontend, connected by a WebSocket channel. Communication to the different devices is made directly through serial communication and whenever necessary, the drivers provided by the vendors or a generic version from the µManager community. CyberSco.Py is installed on computer software that must be equipped with a recent GPU to benefit from U-NET deep learning segmentation of cells. A server can interface several microscopes running CyberSco.Py and be extended with a user manager database and image database management program such as OMERO. The current open-source release of CyberSco.Py can be found on GitHub (https://github.com/Lab513/CyberSco.Py)

### Acquisition, segmentation and tracking

Image acquisition is preceded by an autofocus algorithm that optimizes the quality of the segmentation, as well as the sharpness of the object under scrutiny. Cell segmentation is achieved via a machine-learning algorithm based on the U-NET architecture. Twenty images of cells at different positions and containing different numbers of cells were taken to produce the training set. This dataset was then augmented as it is classicaly done. (see supplementary information). For the main model for yeast segmentation, the neural network was trained on five periods using a GPU NVIDIA GeForce GTX 1080; this training only took five minutes. Predictions with this model are obtained in around 0.2 seconds. Cell tracking is performed using both image correlation in real-time and using a simple proximity relationship between the predicted contours.

## Acknowledgments

The authors would like to thank Pierre Louis Crescitz, Williams Brett and several colleagues and beta testers for their help and critical reading of this manuscript. This work was supported by the European Research Council grant SmartCells (724813) and received supports from grants ANR-11-LABX-0038 and ANR-10-IDEX-0001-02.

## Author Contributions

Code implementation and automation troubleshooting were performed by LC and WB. Experiments were conducted by CC, MLB, AB, SP and DM. JMdM, BS and PH conceived and designed the study. LC, MLB, AB, BS and PH wrote the manuscript.

## Supplementary Information

Supplementary information contains one supplementary text, three figures and three supplementary movies.

