## Supplementary Materials for "CyberSco.Py: open-source software for event-based, conditional microscopy"

### User Documentation

#### 1. Introduction

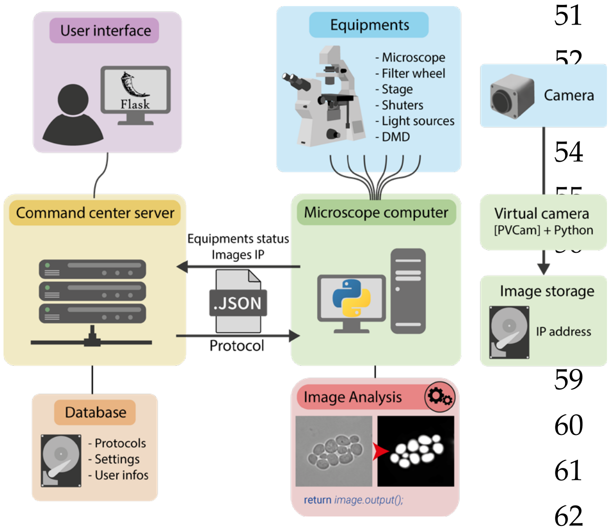

CyberScop.Py is an open-source software that simplifies microscope automation, with an emphasis on conditional and event-based unsupervised timelapse microscopy. The software controls the components of conventional epifluorescent microscope systems (stage positioning, autofocus, illumination settings, camera) and extra devices (Arduino controller, peristaltic pump, syringe pump, serial devices, etc.) and automates their functioning based on the status of the experimental setup using feedback from real-time image analysis. CyberScop.Py enables advanced, event-based experiments to be performed in an unsupervised manner. In contrast with other developments in Python, the focus is on an easy-to-use user interface and predefined scenarios that can be readily used with limited coding expertise. It can be found on [https://github.com/Lab513/CyberSco.Py](https://github.com/Lab513/CyberSco.Py)

In its current version (v. 0.9), the user can either define a MDA from scratch using the graphical user interface or load a predefined scenario—including basic triggers—and adjust relevant parameters. CyberSco.Py critically depends on real-time image analysis, which is currently implemented through the U-NET Convolution Neural Network. We provide trained neural networks; however, as for every deep learning strategy, it is advised to retrain the network on your specific imaging settings.

#### 2. Installation

CyberSco.Py is still in a proof-of-concept stage and requires several installation steps to guarantee that you can use GPU libraries, which are required for Deep Learning Image Analysis to efficiently work Python and related dependencies. This is a critical step. You can find up-to-date instructions on how to install dependent libraries on the GitHub page of the CyberSco.Py project. Do not hesitate to contact us if you need assistance; we will do our best to help you depending on our workload.

##### 2.1 Python dependencies

First, install the Anaconda *Python* distribution platform. Go to the address:
[https://www.anaconda.com/products/individual-d](https://www.anaconda.com/products/individual-d)
Then download and install the version of Anaconda suitable for your OS system.

#### 2.2 NVIDIA GPU libraries installation

Download and install the following libraries:[1]

- CUDA 10.1 > cuda_10.1.243_426.00_win10.exe
  from https://www.filehorse.com/download-nvidia-cuda-toolkit/42676/
- CUDNN 10.1 >> cudnn-10.1-windows10-x64-v7.6.5.32.zip
  visit https://developer.nvidia.com/rdp/cudnn-archive and from the unfolded list, select *Download cuDNN v7.6.5 (November 5th, 2019), for CUDA 10.1*

You then need to copy three sets of files from CUDNN to CUDA

- `<cuDNN directory>\cuda\bin\*.dll` >> `C:\Program Files\NVIDIA GPU Computing Toolkit\CUDA\vxx.x\bin`
- `<cuDNN directory>\cuda\include\*.h` >> `C:\Program Files\NVIDIA GPU Computing Toolkit\CUDA\vxx.x\include`
- `<cuDNN directory>\cuda\lib\x64\*.lib` >> `C:\Program Files\NVIDIA GPU Computing Toolkit\CUDA\vxx.x\lib\x64`

Now, check you have the following two paths in the “Environment Variables” list:

```
C:\Program Files\NVIDIA GPU Computing Toolkit\CUDA\vxx.x\bin
C:\Program Files\NVIDIA GPU Computing Toolkit\CUDA\vxx.x\libnvvp
```

To do so, open the Start Search, type in “env”, and choose “Edit the system environment variables”. Click the “Environment Variables…” button. The two paths above should be found by scrolling through the “System variables panel”.

You can check that your CUDA and CUDNN are installed correctly by opening “Control Panel”, click “System and Security”, and then click "Device Manager." Open the "Display Adapters" section, double click on the name of your graphics card and then look for the information under "Device status." This area will typically say "This device is working properly" if it is OK.

#### 2.3 Coolled driver installation

We rely on the driver proposed by the µManager community. Information on the driver installation and serial virtual ports can be found at: https://micro-manager.org/wiki/CoolLED. More generally, the µManager community provides many drivers that can be used to control your own microscope through µManager, and also through Python, either directly or using the PycroManager library (https://pycro-manager.readthedocs.io/en/latest/)

#### 2.4. Camera installation

At present, we are using a Zyla (from Andor) and an Evolv512 (from Roper Scientific).

<u>Installation of the Zyla camera</u>

[1] We are using a NVIDIA GPU.(GTX 1080 or Quadro RTX4000). If you are using a different GPU, you should adapt this step.

in the folder driver/pyAndorSDK3, open a console and run :

```
> python -m pip install .
```

Do not forget the dot after "install" !

Installation of Evolv512(Pyvcam)

All of the information can be found at https://github.com/Photometrics/PyVCAM

In brief:

a) Download and install the drivers for the camera from https://www.photometrics.com/support/software-and-drivers#software

b) Download the GitHub folder from the address above.

c) Unzip and open a console in this folder

d) Write and execute `> python setup.py install`

#### 2.5 Microfluidics installation

The Arduino used to control the microfluidic valves uses the program *ValvesArduinoScript.ino,* which can be found in the CyberSco.py folder at the address:

```
> CyberSco.py/drivers/ValvesArduinoScript
```

The software for interacting with the Arduino can be found at https://www.arduino.cc/en/software

#### 2.6 Port settings

The correct port settings are required to make sure that the devices can be found properly by CyberSco.Py. We are working on an automated device/port discovery for the next version. For now, we proceed manually by searching Windows Device manager to find which port each device is associated with (once it has been plugged and installed). For example, in our configuration we have:

- Olympus IX81: COM1, 19200 bauds
- PRIOR stage: COM4, 19200 bauds
- COOLLED : COM5, 38400 bauds
- Arduino: COM8, 9600 bauds
- XCite: COM9, 9600 bauds

A port configuration yaml file that contains all of the serial communication parameters exists for each device. These configuration files have to be placed in :

```
> CyberSco.py/modules/settings/ports/
```

For example, the "coolled.yaml" file for the COOLLED device is:

```
# port for Coolled

port : COM5
baudrate : 38400
bytesize : EIGHTBITS
parity : PARITY_NONE
stopbits : STOPBITS_ONE
xonxoff :  False
```

Changing the ports in Windows :

1. Go to Windows Device manager > Multi-port serial adapters.
2. Select the adapter and right click to open the menu.
3. Click on the Properties link.
4. Open the Ports Configuration tab.
5. Click on the Port Setting button.
6. Select the Port Number and click OK.
7. Click OK to apply the changes.

#### 2.7 CyberSco.Py installation

You first need to retrieve the Python code from the GitHub repository. Download and unzip the Zip file. Go into the CyberSco.py folder, open a console window, then write and execute:

```
> python setup.py install
```

If the installation works, you should see a new icon on your desktop.

#### 2.8 Machine Learning Models

The models produced with TensorFlow must be placed in the folder
`> CyberSco.py/models`
Then, you need to create a file *models.yaml* in
`> CyberSco.py/modules/settings`
This file contains the correspondences between the models and their shortcut.
Two other configuration yaml files must also be provided in the same folder.
The first is *curr_models.yaml*, which contains the shortcut for the main segmentation model in use.
The second is *event_model.yaml*, which contains the shortcut for the model devoted to event detection.

#### 3. Launch CyberSco.py

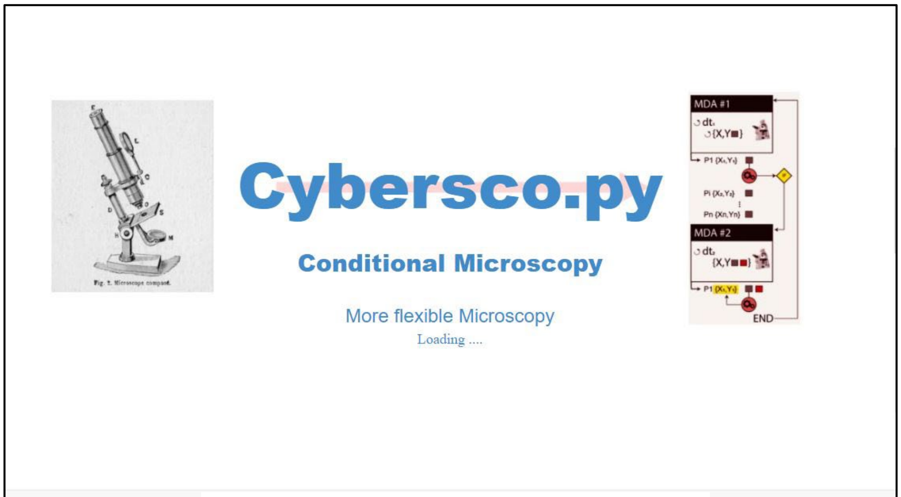

Figure 1 Starting window of CyberSco.Py

Turn on your microscopes and devices. Then click on the icon on the Desktop. You can alternatively open an Anaconda Terminal from the folder CyberSco.py and run the following command:

```
> python -m interface.run
```

The program opens a window, and loads the different components and tests the connection with the devices, which takes about 30 seconds to one minute. Once it is ready, a *Begin* button appears. Just click on it.

#### 4. Main interface window

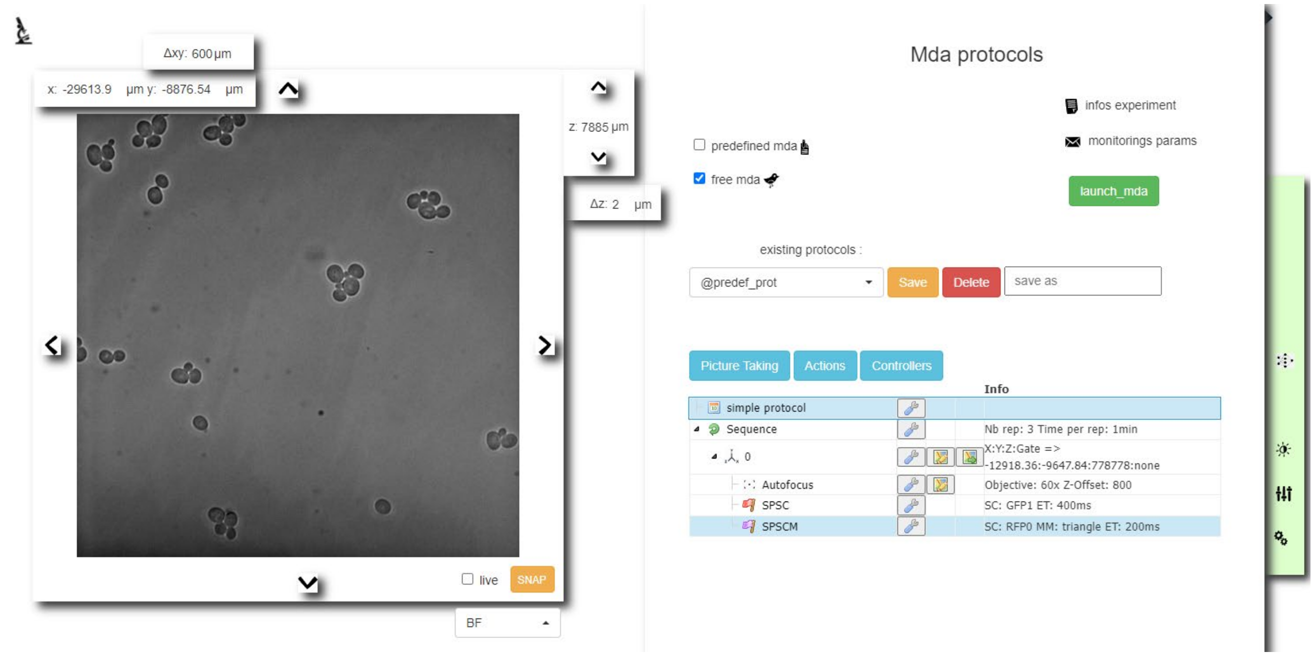

The window shows the field of view of the camera and the buttons to interact with the microscope. A green vertical menu bar with four buttons is on the right side of the interface. From top to bottom we find: the button for opening the machine learning monitoring panel, the button for the Settings Channels, the button for performing a panoramic view of the chip, and the button for opening the panel that contains information about the devices currently connected and the GPU.

**Moving the stage in X-Y directions**

The X-Y coordinates of the current position appear at the top of the window. Four arrows on the edges of the window permit the field of view to be displaced along the four cardinal directions. The position can also be modified by directly entering the coordinates.

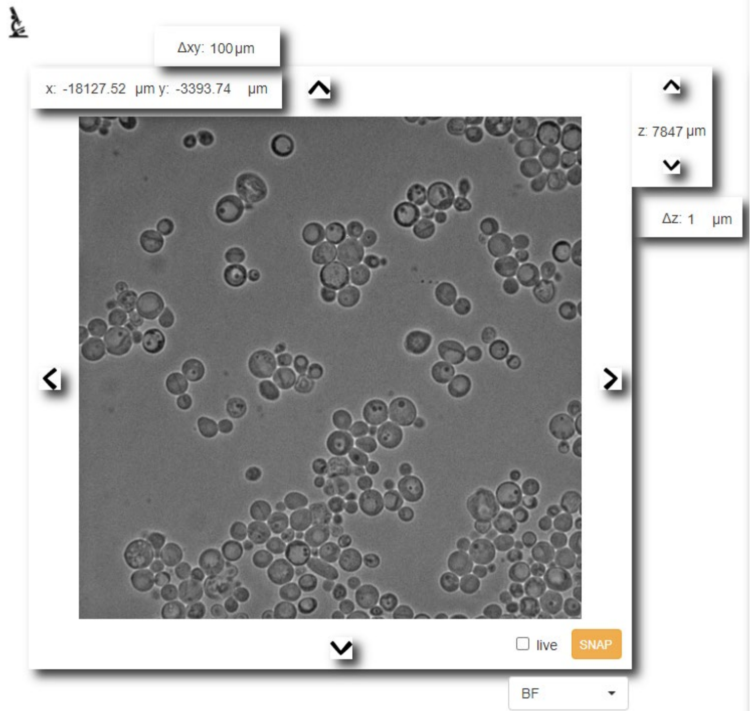

**Changing the focal position**

The current Z-position of the microscope is displayed between the vertical arrows on the left. The Z-position can be changed by directly entering the z-value or by clicking on the arrow buttons. The z-step can also be modified by entering its value or by clicking on the up-down arrow buttons.

**Taking a snapshot**

The SNAP button (orange button) for taking a snapshot under different light conditions can be found at the bottom-right of the panel that displays the current microscope view. Before clicking on SNAP, the light conditions must be selected using the selector under the button, which proposes different *settings channels* registered previously by the user. Alternatively, the user can click on the checkbox on the left of the orange button to permanently activate the selected setting channels. Unchecking this box deactivates the illumination. The exposure time can be modified by right clicking on the snap button and changing the value in milliseconds.

**Overlaying BF and fluorescence images**

After acquisition of a snapshot fluorescence image, the *blue slider* under the overlaid snapshot can be used to set the opacity of the fluorescent image above the BF view of the field of view. The cross on the top right corner of the snapshot is used to close the snapshot view and return to the current BF view.

**Settings channels**

This panel is used to define the illumination settings used in live mode or an acquisition timelapse. To open the settings panel, click on 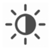

The user needs to click on the checkboxes of the objects in the settings channel to be used (here, we have three light sources: BF, Xcite, Coolled). To specify the values for each of those objects, give a name to the settings channels and save it. The setting channels will then appear in the selector for the snapshot, as well as in the tree for making the MDAs. For the Xcite device, a wavelength filter can be associated by selecting a number in the select box.

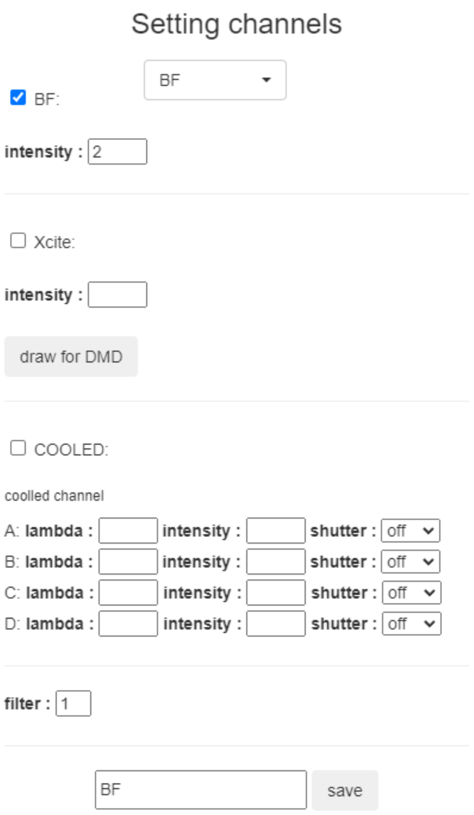

Each settings channel is saved in the folder *interface\settings\settings_channels* as a *yaml file*

```yaml
name_set_chan: RFP0
BF: '100'
COOL:
  A:
    lamb: '365'
    intens: '1'
    shut: 'off'
  B:
    lamb: '460'
    intens: '3'
    shut: 'off'
  C:
    lamb: '550'
    intens: '4'
    shut: 'on'
  D:
    lamb: '635'
    intens: '1'
    shut: 'off'
```

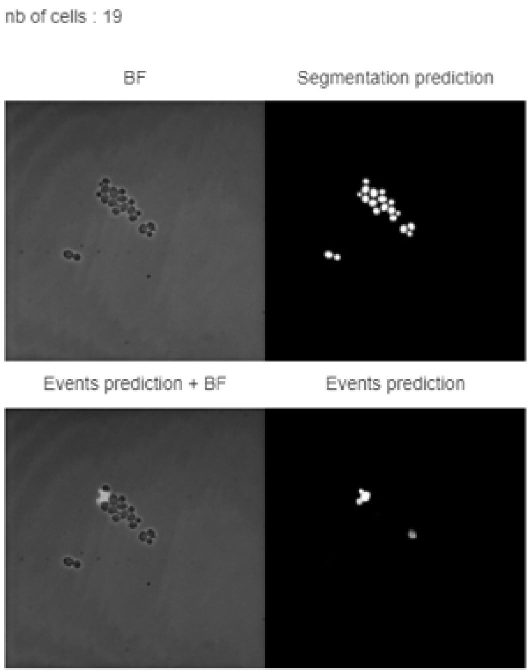

**Segmentation monitoring**

When setting-up an experiment, this panel is useful to assess the quality of the segmentation in real-time before launching the time-lapse. To open it, click on 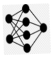

This tool also allows the user to test the Machine Learning model's robustness in live mode (i.e., how the model behaves with the current field of view, changing the position of the *z*-axis, etc...). The first line shows the result of the segmentation for the main segmentation model. The second line shows the segmentation for a supplementary loaded model. The main model is devoted to cell segmentation, and was the model used in all of the experiments reported in this study. The supplementary model is an optional model that can be used to detect specific events, like *budding cells* as shown in the picture above.

#### Settings and tools

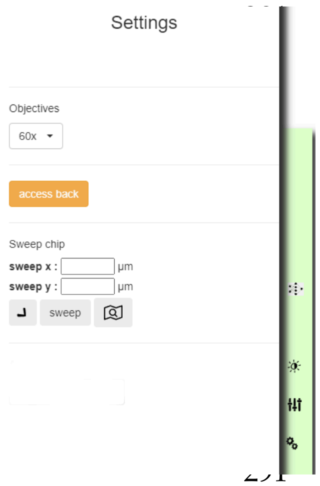

To access this panel, click on

##### Set the objective

This panel allows the user to change the objective, and also displays the current objective in place.

##### Hand-back function

When the CyberSco.Py interface is open, the knob/wheel that controls the focus of the microscope will be inactive. Clicking on the "*hand back*" button allows the user to adjust the knob to tune the focus manually.

#### Sweep chip tool

While preparing an experiment, it can be useful to have a wider overview of the sample than the field of view set by the objective. The tool *"sweep chip"* allows the user to acquire multiple images across the sample and displays the resulting stitched tiles. You can choose the size of the area to be scanned. When the overview is displayed, you can click on the window where you want the field of view to be centered.

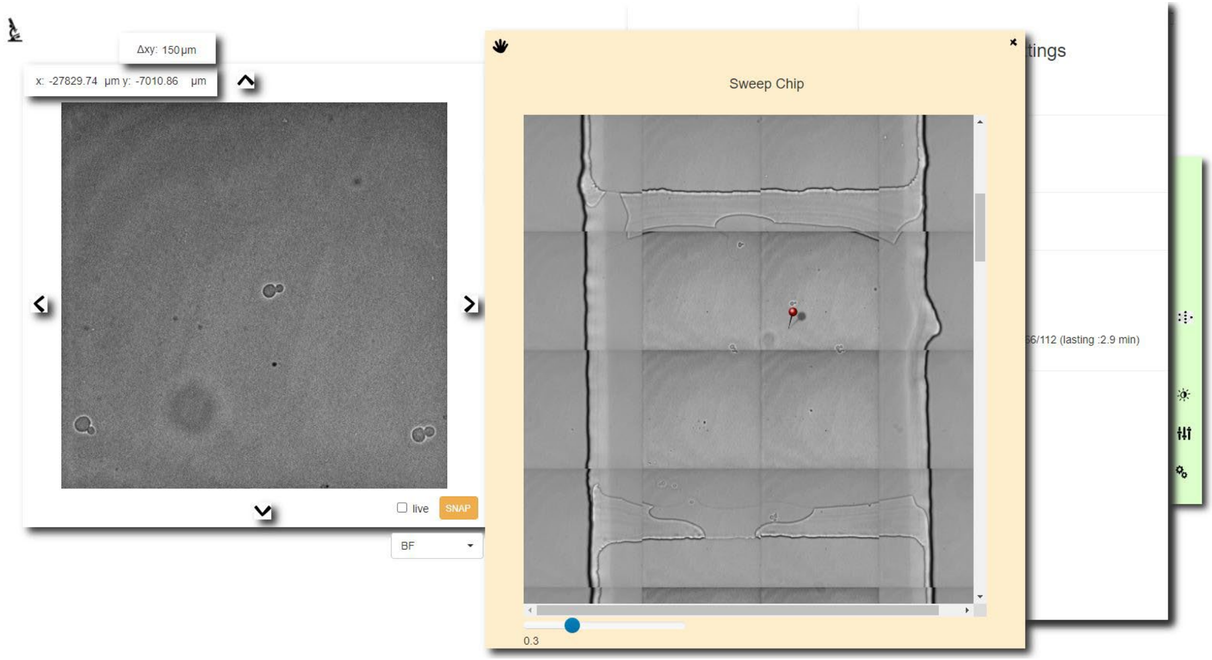

**Connected devices, GPU information and TensorFlow version**

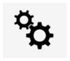

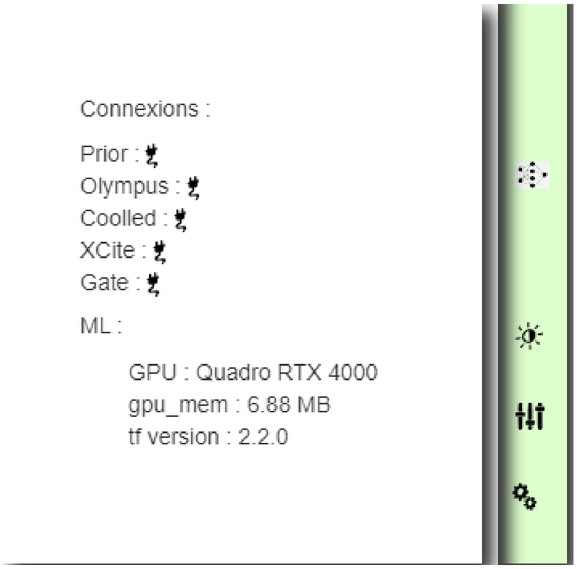

This panel indicates the devices that are correctly connected (showing a plug icon if a device is connected correctly), which model of GPU is used and its memory, and which version of TensorFlow will run the experiments.

#### 5. Defining and using a MDA experiment

The *MDA protocols* panel allows the user to choose between two kinds of MDA experiments: *free MDA* and *predefined MDA.* Below, we explain how the user creates or sets the parameters of the experiment for each case.

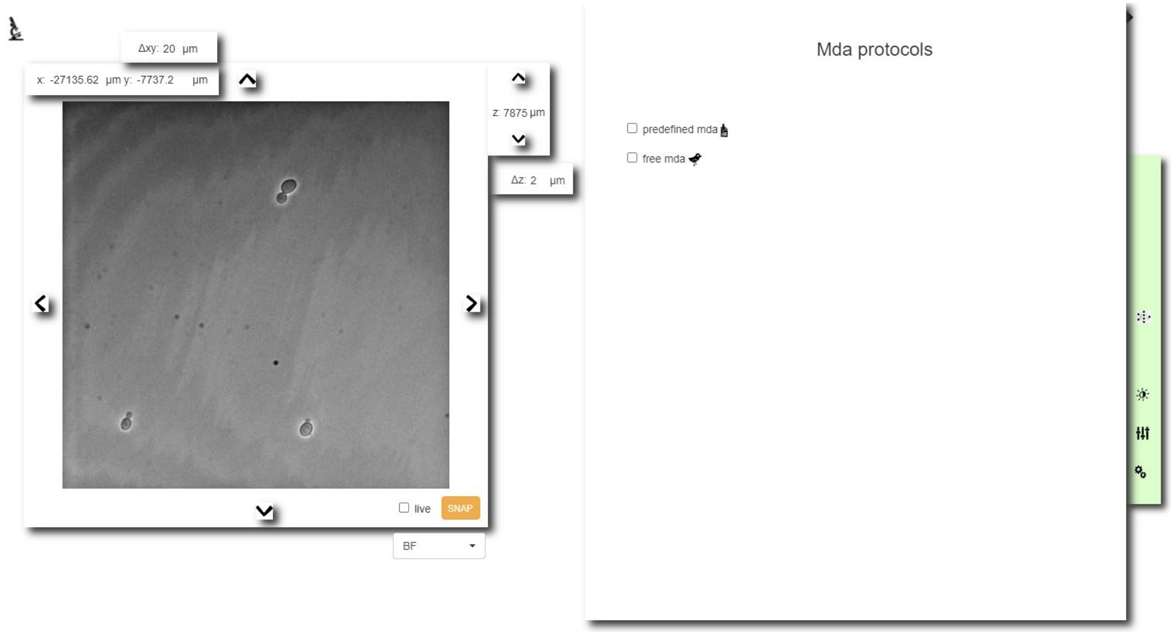

**Free MDA experiment**

Select the option *free MDA* in the MDA protocols panel. A tree for creating, saving, modifying and duplicating MDAs will appear.

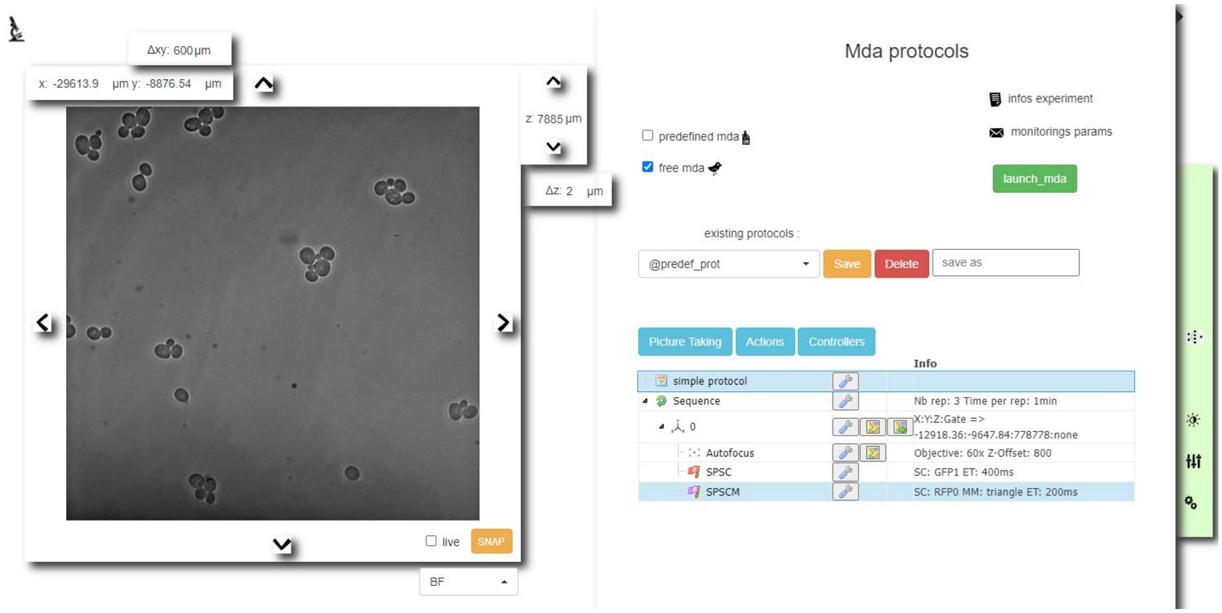

The tree graphically encapsulates the serial instructions for the microscope with associated loops. The most basic structure is a loop over a position, which contains the instructions for *taking a picture in brightfield mode*. The user can duplicate, erase or modify part of the tree or create new loops and instructions from scratch; it is also possible to create nested loops. To add new elementary blocks, just drag and drop the blocks from the three selection menus (the three blue selectors just above the tree) into the tree. You can modify the hierarchical position of the blocks using *ctrl+arrow*.

**Changing the parameters of each action**

To modify the parameters of each element in the tree, click on the corresponding wrench icon and a window will open. You can change and save new sets of parameters.

**Predefined MDA experiment**

Select the option *predefined MDA* in the panel *MDA protocols*. A panel with suggestions of predefined experiments will appear on the left. Choose the predefined MDA you require.

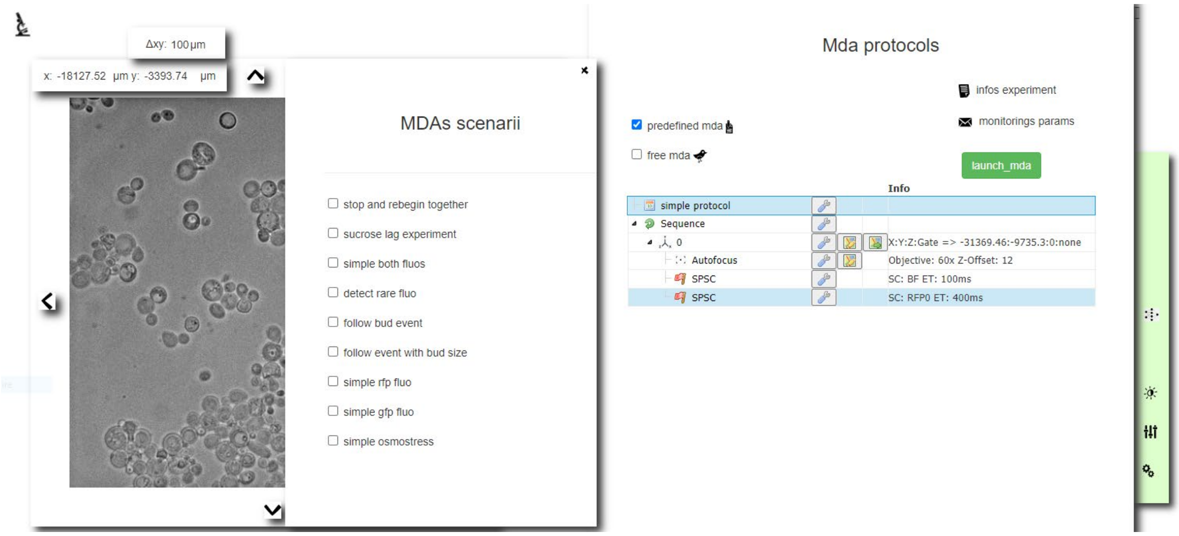

When *predefined MDA* is chosen, a tree will also appear at the bottom of the panel, which permits the user to define the positions on which the *predefined MDA* will run. The tree functionalities are the same as for creation of the free MDA protocol.

**Storing additional experimental information**

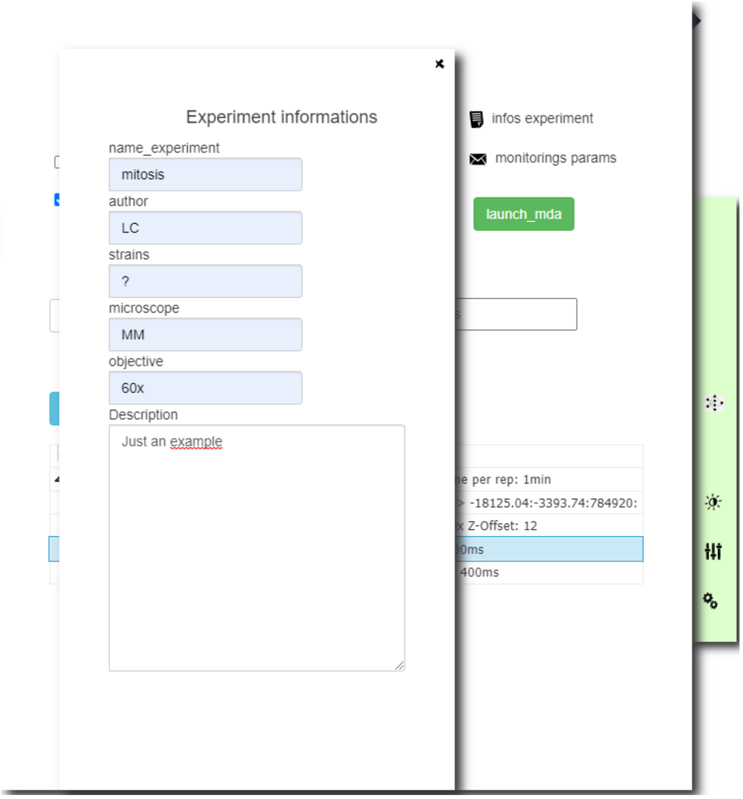

For each experiment, the user can enter information like the *name of the experiment, authors' names, information about the biological system*, the *microscope* in use, the *objective* and supplementary *free descriptions and remarks*. The resulting file is saved in the results folder as *infos.txt*.

#### 6. MDA results

The experimental results, the monitoring images and parameters, and information about the experiment are saved by default in a folder named *mda_temp* in the CyberSco.py folder. A log.dat file is also produced to register, in real-time, the events the user wanted to follow during the experiment, and to trace eventual bugs.

#### Advanced: Creation of a new preprogrammed MDA

Each preprogrammed experiment is stored as a unique Python file. These files must be placed in

```
> CyberSco.py/modules/predef/plugins/
```

The program consists of a python class with the name of the experiment, which is inherited from the MDA class. This class contains an *__init__* block (containing in the comments the text that will appear as tooltip in the interface for the plugin description) and a *define* block, which contains the instructions for the MDA written serially followed by the *self.launch_loop()* instruction.
There are two blocks to initialize the experiment: *init_on_positions()* and *init_conditions()*.
The last block is the *check_conditions()* block, which will be executed after each acquisition.
Below, we show an example of the code for a plugin :

```
from datetime import datetime
from modules.mda import MDA

class PROTOCOL_SUCROSE(MDA):
    '''
    Comments about the plugin
    '''
    def __init__(self, ldevices=None):
        '''
        name : experiment name
        description : experiment description
        '''
        MDA.__init__(self, ldevices)

    def define(self, debug=[0]):
        '''
        '''
        self.refocus()                          # add refocusing
        self.take_pic()                         # add take BF pic
        self.analyse_pic()                      # analyse the pic
        self.cond = 'sucrose'                   # apply the conditions 2,
sucrose
        self.launch_loop()                      # Loop

    def init_on_positions(self):
        '''
        Initialize the protocol
        '''

    def init_conditions(self):
        '''
```

```
        Setup parameters and initial conditions
        '''

    def check_conditions(self, rep):
        '''
        At a given threshold trigger the sucrose
        '''
        for pos in self.list_pos:
            if pos.nb_cells > pos.thresh_cells and not pos.switched :
                if pos.num_gate:
                    pos.switched = True
                    self.gates_switched += [ pos.num_gate ]
                    self.ga.set_pos_indices( self.gates_switched, 1 )
```

### List of connected devices

| Name | Function |
|---|---|
| Olympus IX81 | Fully automated microscope (including ZDC focus, filter wheel turret, motorized focus, shutters, ...) |
| pE-4000 Cooled | Fluorescent illumination |
| Prior ProScan III | Stage XY displacement |
| Photometrix Evolve512 | Camera |
| ISMATEC IPC ISM932D | Peristaltic pump |
| Arduino | Valve control |
| XCite | Fluorescent illumination |

### Image analysis method to extract information from cells

The experiments presented in this work rely on the utilization of two segmentation models within the U-NET architecture. Training was performed using a Nvidia GeForce GTX 1080 GPU card with TensorFlow 2.2.0 and with data augmentation with rotation (six angles), flip operation, noise, colors, and contrast.

#### 1. Segmentation

The experiments in this paper rely on two segmentation models; the first allows the user to segment yeast cells at 20x and the second, at 60x. The training sets were obtained from RFP images of nuclear-tagged yeast cells. We used the OTSU thresholding algorithm to automatically create cell masks.

The first model (20x) was trained over 15 epochs (in 15 minutes) using a training set of 20 pictures. The figure below shows an example of segmentation on a 20x BF field image with the first model.

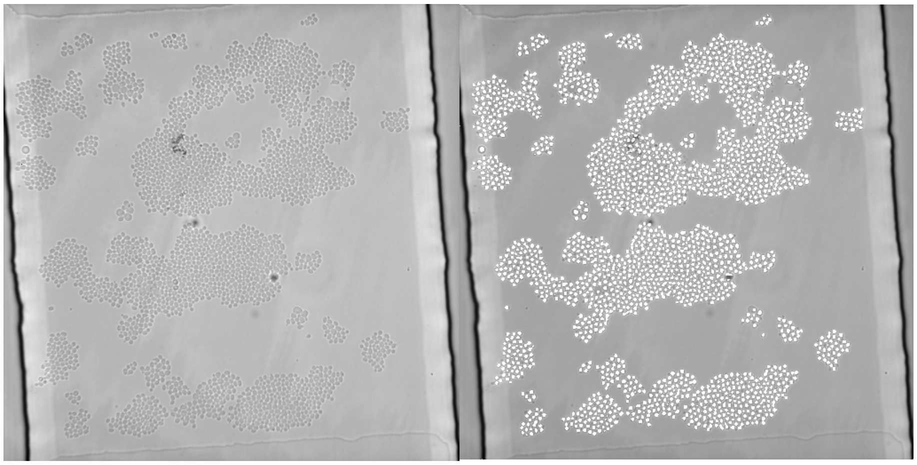

The second model (60x) was trained over 5 epochs (in 5 minutes) using a training set of 20 pictures. Below, we show an example of segmentation on a 60x BF field image with the second model.

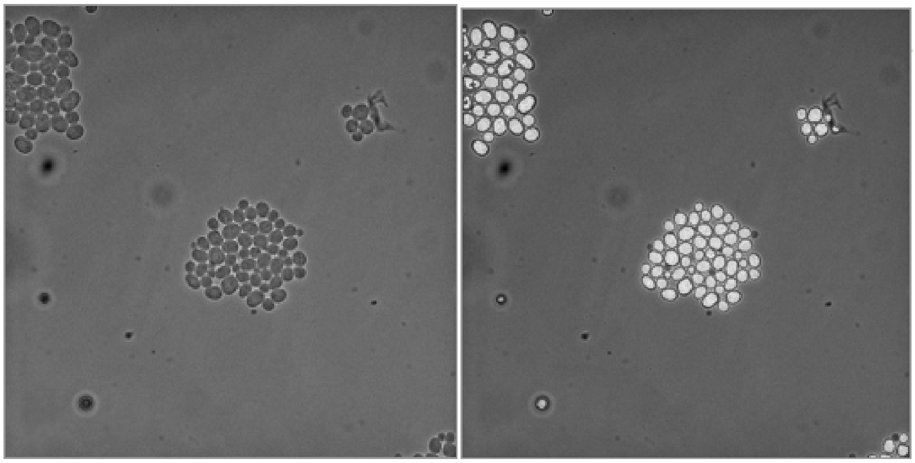

The two training sets used to build the segmentation models are available on the GitHub repository of the project. The data were augmented and split into training and validation groups using the scikitlearn *train_test_split* method in a ratio of 95% of the images for training and the remaining 5% for validation.

#### 2. Tracking

Tracking is easily achieved by linking the cells that are the closest from one image to the following image. For each experiment, the tracking pictures are stored in the folder *monitorings/tracking.*

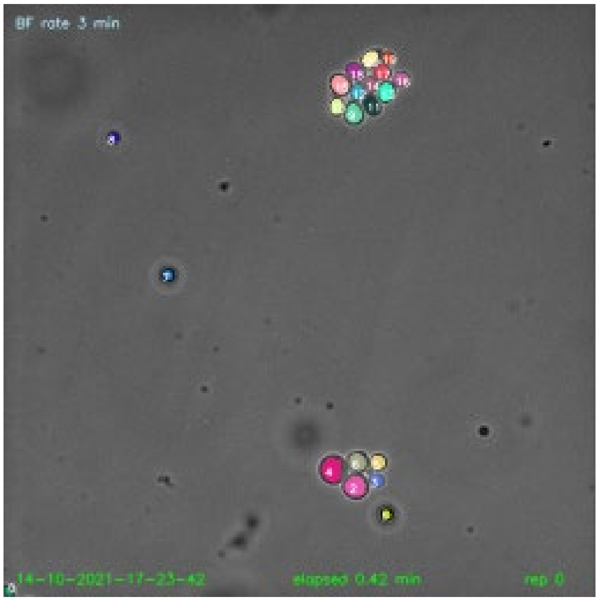

### Supplementary Table 1

| Strain name | HO locus | HIS locus | HOG1 locus | Nuclear marker | Background |
|---|---|---|---|---|---|
| yPH428 | | | | | BY4741 |
| yPH449 | | EL222-HIS3 | | HTB2::mApple-Kan | BY4741 |
| yPH459 | $P_{C120}$-Venus | EL222-HIS3 | | HTB2::mApple-Kan | BY4741 |
| yPH15 | | | HOG1::GFP-HIS3 | HTB2::mCherry-URA3 | BY4741 |

Supplementary Table T1 – List and genotypes of yeast strains used in the study. yPH428 was used for Fig. 4 and Fig. 5. yPH449 was used for Fig. 2.A, Fig. 4 and Fig. 6. yPH459 was used for Fig. 2.B. yPH15 was used for Fig. 3.

### Supplementary Figure S1

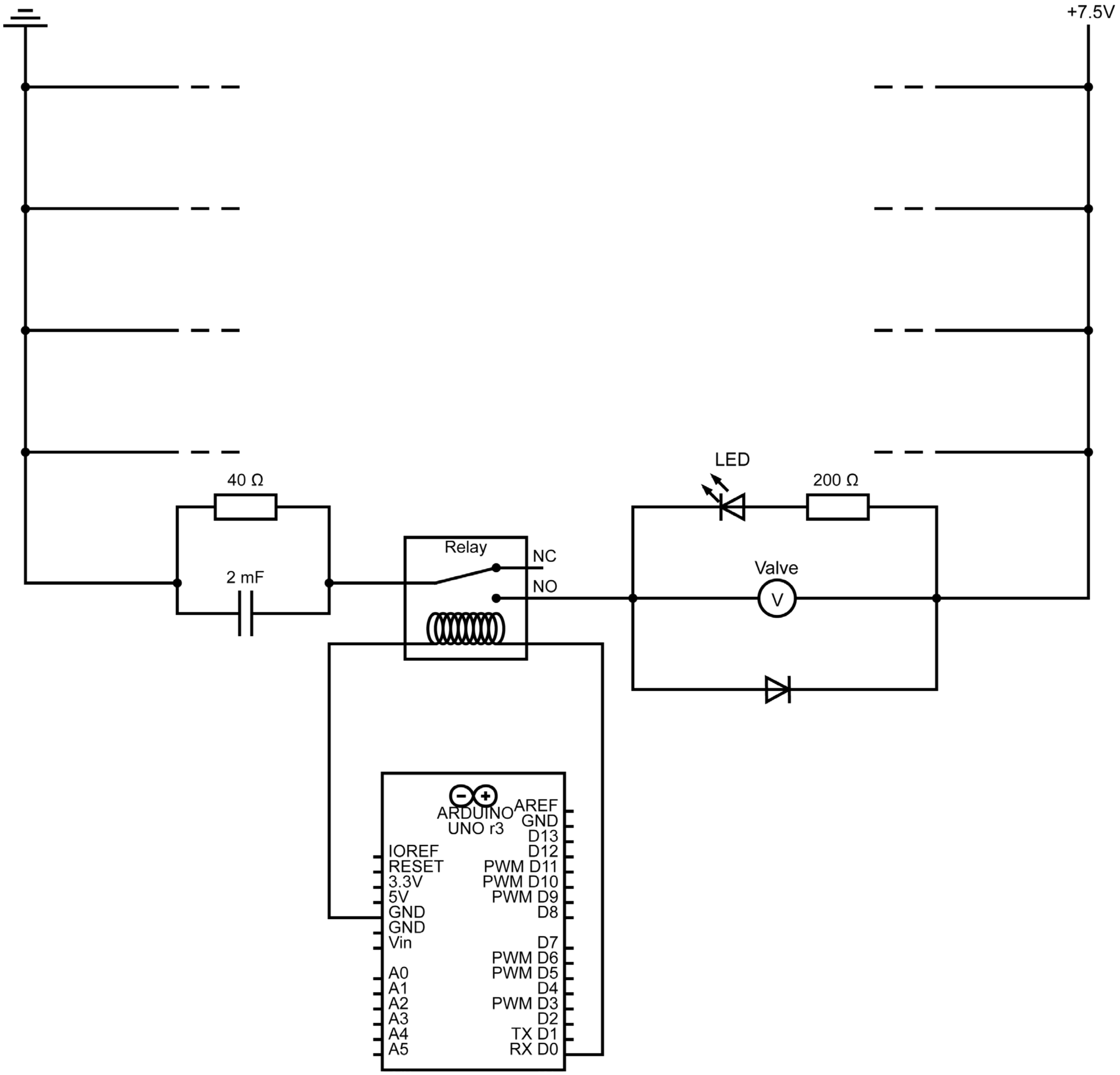

**Supplementary Figure S1** – Valve control electronic scheme. The capacitor part of the circuit introduces a spike of 7 V when switched to ON, then applies a lower voltage (3.5 V) when the solenoid valve is maintained in the ON position. The presence of the capacitor makes the switching more reliable compared to a single resistor. We used LHDA0531115H Lee company valves. We control the Arduino using serial communication, updating the state of each valve (either 0 or 1). This system can be controlled independently of CyberSco.Py using a simple Jupyter notebook..

### Supplementary Figure S2

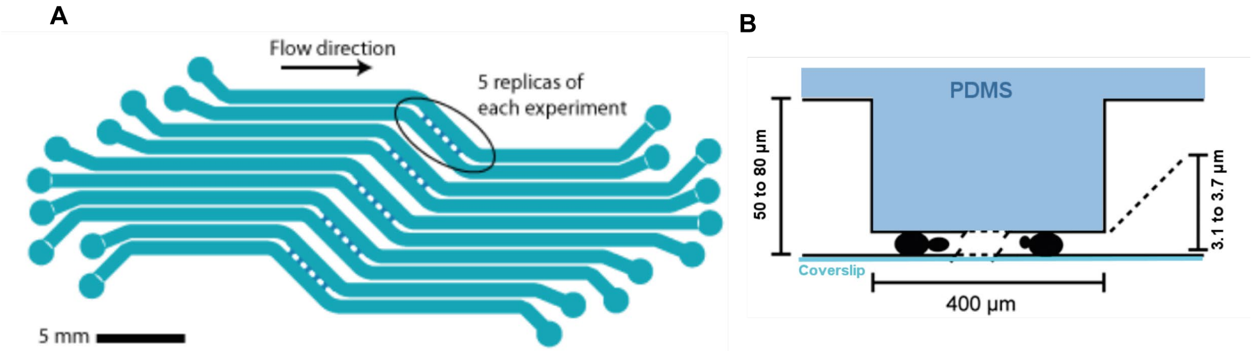

**Supplementary Figure S2** – Microfluidic chip design. (A) Top view showing that five experiments can be run in parallel, with five replicates for each. (B) Side view. Yeast cells are sandwiched between glass and PDMS, which ensures the cells grow as a monolayer that is amenable to live cell segmentation and tracking. This design is regularly used by our team for long-term time-lapse microscopy of yeast cells under fluctuating environmental conditions. Specific designs and operating instructions are available upon request.

### Supplementary Figure S3

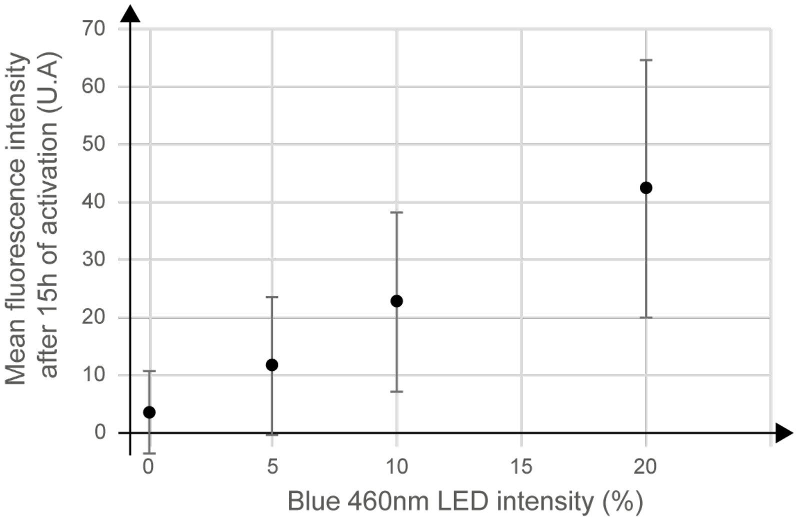

**Supplementary Figure S3** – Light dose response of the optogenetic promoter $P_{C120}$, constructed from one experiment, as shown in Figure 2 of the main text. Fluorescent intensity was measured over the whole chamber filled with yeast. Error bars represent ± the standard deviation of pixel intensity.

#### Supplementary Movies SM1,SM2,SM3

These movies demonstrate how to use the user interface to create simple and advanced time lapse with CyberSco.py.
